# Loss of EIF4G2 Mediates Aggressiveness in Distinct Human Endometrial Cancer Subpopulations with Poorer Survival Outcome in Patients

**DOI:** 10.1101/2023.09.14.557672

**Authors:** Sara Meril, Maya Muhlbauer Avni, Chen Lior, Marcela Bahlsen, Tsviya Olender, Alon Savidor, Judit Krausz, Hila Belhanes Peled, Hila Birisi, Nofar David, Shani Bialik, Ruth Scherz-Shouval, Yehuda Ben David, Adi Kimchi

## Abstract

The non-canonical translation initiation factor EIF4G2 plays essential roles in embryonic development and differentiation, and contributes to the cellular stress response via translation of selective mRNA cohorts. Currently there is limited and conflicting information regarding the potential involvement of EIF4G2 in cancer development and progression. Endometrial cancer (EC) is the most pervasive gynecological cancer in the developed world, with increasing incidence every year. High grade ECs are largely refractory to conventional treatments, presenting poor survival rates and lacking suitable prognostic markers. Here we assayed a cohort of 280 EC patients across different types, grades, and stages, and found that low EIF4G2 expression highly correlated with poor overall and recurrence free survival in Grade 2 EC patients, monitored over a period of up to 12 years. To establish a causative connection between low EIF4G2 expression and cancer progression, we analyzed in parallel two independent human EC cell lines and demonstrated that stable EIF4G2 knock-down resulted in increased resistance to conventional therapies. Depletion of EIF4G2 also increased the prevalence of molecular markers for aggressive cell subsets, and altered their transcriptional and proteomic landscapes. Prominent among the proteins with decreased abundance were Kinesin-1 motor proteins KIF5B and KLC1, 2, 3. Multiplexed imaging of the tumors from this EC patient cohort showed a correlation between decreased protein expression of either KIF5B or KLC1, and poor survival in patients of certain grades and stages. The findings herein reveal potential novel biomarkers for Grade 2 EC with potential ramifications for patient stratification and therapeutic interventions.

**Significance:** Decreased EIF4G2 protein results in increased drug resistance of aggressive sub-populations of endometrial cancer cells, is associated with poor patient survival, and may serve as a novel prognosis marker for endometrial cancer.

## Introduction

Enhanced protein translation is a characteristic of cancer cells, which have increased metabolic needs stemming from their rapid cell division and growth. As such, several canonical translation initiation factors are known oncogenes (1). Recently, the non-canonical translation initiation factor EIF4G2 (also known as DAP5/p97/Nat1 (2–5)) has attracted interest in the cancer field. A scaffold protein that recruits the 40S ribosome to the 5’ end of the mRNA, EIF4G2 drives translation independently of the mRNA’s 5’cap structure and/or canonical cap-binding proteins through various alternative mechanisms (6–10). Importantly, EIF4G2 is necessary for embryonic development at early stages (11–13) and differentiation of both human and mouse embryonic stem cells (hESC, mESC) (14,15). Among EIF4G2’s mRNA targets are pro- and anti-apoptotic proteins that are specifically translated during apoptosis (16–18) and mitosis (19), an N-terminally truncated isoform of p53 that is translated during cell stress (20), and epigenetic modulators such as HMGN3 and KMTD2 that are essential for hESC differentiation (15,21). These indicate an important role for EIF4G2 in the translation of factors critical for cell fate decisions in somatic and embryonic stem cells, and it is not surprising that it has been linked to cancer.

Paradoxically, a limited number of studies have attributed both oncogenic and tumor-suppressive capabilities to EIF4G2. In triple negative metastatic breast cancer, EIF4G2 was shown to mediate non-canonical translation of targets involved in cell migration, the epithelial-mesenchymal transition (EMT) and invasion, and its overexpression correlated with and promoted tumor metastasis (22). In the same vein, a correlative study of EIF4G2 in gastric cancer showed that increased *EIF4G2* mRNA expression was associated with poor prognosis (23). On the other hand, *EIF4G2* expression was reduced in bladder cancer, correlating with tumor de-differentiation and invasiveness (24). Our recent analysis of the TCGA database demonstrated a significant reduction in *EIF4G2* mRNA expression in 7/24 primary tumor types compared to healthy tissue, as opposed to 2 tumor types (i.e., glioblastoma multiforme and cholangiocarcinoma) that showed significant increased mRNA expression (BioRxiv 2023.08.22.554280). Moreover, we identified deleterious mutations (out-of-frame deletions, insertions and nonsense mutations) and significantly occurring somatic missense mutations in *EIF4G2* in various cancers, several of which were proven to be loss-of-function. Thus, it appears that EIF4G2’s contribution to cancer varies according to the type and/or stage of cancer. Notably, endometrial cancer was prominent among the cancers showing reduced *EIF4G2* expression and the presence of somatic mutations, representing 14% of all occurrences of the mutations among all examined cancer types (BioRxiv 2023.08.22.554280).

Endometrial cancer (EC) is the most common gynecological cancer in developed countries, with increasing incidence in recent years due to the rising prevalence of obesity, a main risk factor for the disease (25). EC staging (1–4) according to the International Federation of Gynecology and Obstetrics (FIGO) indicates the degree of invasiveness, with stage 1 limited to the corpus uteri and later stages progressively involving increased invasiveness and distant metastases. FIGO also grades tumors on a scale of 1-3 according to the relative proportions of the glandular and solid-tumor components. Grade 1 is well differentiated, Grade 2 is considered moderately differentiated, and Grade 3 is poorly differentiated, with a solid-tumor component less than 6%, 6-50%, and more than 50%, respectively. Buckham’s dualistic classification combines several factors, among them histological sub-type, grade, and hormone dependency into Type-1 and Type-2 EC. Type-1 hormone-dependent endometrioid cancer affects approximately 80% of patients, mostly classified as low grade (Grades 1 and 2 endometrioid adenocarcinoma) with favorable prognosis (5-year overall survival (OS) rate of 85%). Type-2 EC, diagnosed in the remaining 20% of patients, is characterized by hormone-independent, high-grade (Grade 3) endometrioid adenocarcinomas, serous clear cell, undifferentiated and carcinosarcomas tumors, with a higher risk of metastasis and a poorer prognosis (5-year OS rate of ∼55%) (26).

Standard treatment for EC is surgical resection, and depending on tumor stage and grade is often accompanied by adjuvant therapy (26). The most common adjuvant therapies are radiation and chemotherapy. While survival rates for patients with low grade, early detected tumors can be as high as 95%, high grade and recurrent tumors are more refractory to treatment, and overall survival rates drop to 15-17% (27). Thus, there is still a pressing need for new and advanced therapies and prognostic markers for the successful treatment of EC. One promising approach involves specifically targeting aggressive subpopulations within EC tumors with stem-like characteristics that can be identified by expression of certain markers such as CD133, CD44 and ALDH1A1 (27). These cancer sub-populations, which have been identified in reproductive cancers such as breast and endometrial cancers and various other cancers types, including myeloid leukemia, melanoma, prostate and colon cancer, have been shown to acquire increased metastatic capacity and invasiveness, increased resistance to chemotherapies and radiotherapies, and increased tumorgenicity. As their presence is presumed to confer tumor aggressiveness, strategies to reduce their growth or induce their differentiation have been recently promoted (28).

Here we assessed the contribution of EIF4G2 to EC by examining its protein expression in a cohort of 280 EC patients of varying types, grades and stages. Low expression of EIF4G2 was associated with decreased overall survival of patients with Grade 2 EC. To evaluate whether this association was causative, EIF4G2 was depleted from HEC-1A and RL95-2 EC cell lines, which showed increased resistance to Taxol and radiation treatment, and enrichment for cells with high expression of CD133 and CD44. Moreover, EIF4G2 knock-down (KD) specifically altered the transcriptomic and proteomic signatures of CD133 positive cells, which were particularly resistant to chemo- and radiation therapy. Among the proteins with decreased abundance were direct translation targets of EIF4G2, such as the kinesin-1 motor protein; conversely, DNA damage repair proteins increased in abundance as an indirect response to EIF4G2 KD. The clinical significance of these observations was assessed by multiplexed imaging of the tumors from the EC patient cohort, which showed a correlation between decreased protein expression of kinesin-1 proteins KIF5B and KLC1, and decreased overall survival rates in patients with more advanced EC. Our results show the prognostic potential of EIF4G2 and its potential protein targets.

## Results

### Low protein expression of EIF4G2 correlates with poor prognosis in endometrial cancer patients

The clinical significance of changes in expression of EIF4G2 in EC was assessed in a cohort of 280 endometrial cancer patients collected and followed for up to 12 years post-surgery. Details on patient data, tumor histology, treatments received, metastases, survival and recurrence are shown in Supplemental Table 1. All underwent surgical resection of the uterus. Some patients received adjuvant treatment following surgery, such as radiation therapy (brachytherapy or external radiation) and/or chemotherapy (carboplatin, Taxol). Kaplan-Meier analysis of overall survival (OS) of the samples recapitulated the expected patient outcomes based on type, grade, and stage of EC. Type-2 EC showed significantly lower survival compared to Type-1 EC (Fig. 1A). Similarly, Grade 3 tumors had lower survival rates compared to lower grade tumors (Fig. 1B). Survival also correlated with the tumor stage, with the more advanced stages 3 and 4, associated with increased metastasis, showing the worst OS outcomes (Fig. 1C).

**Fig. 1.**
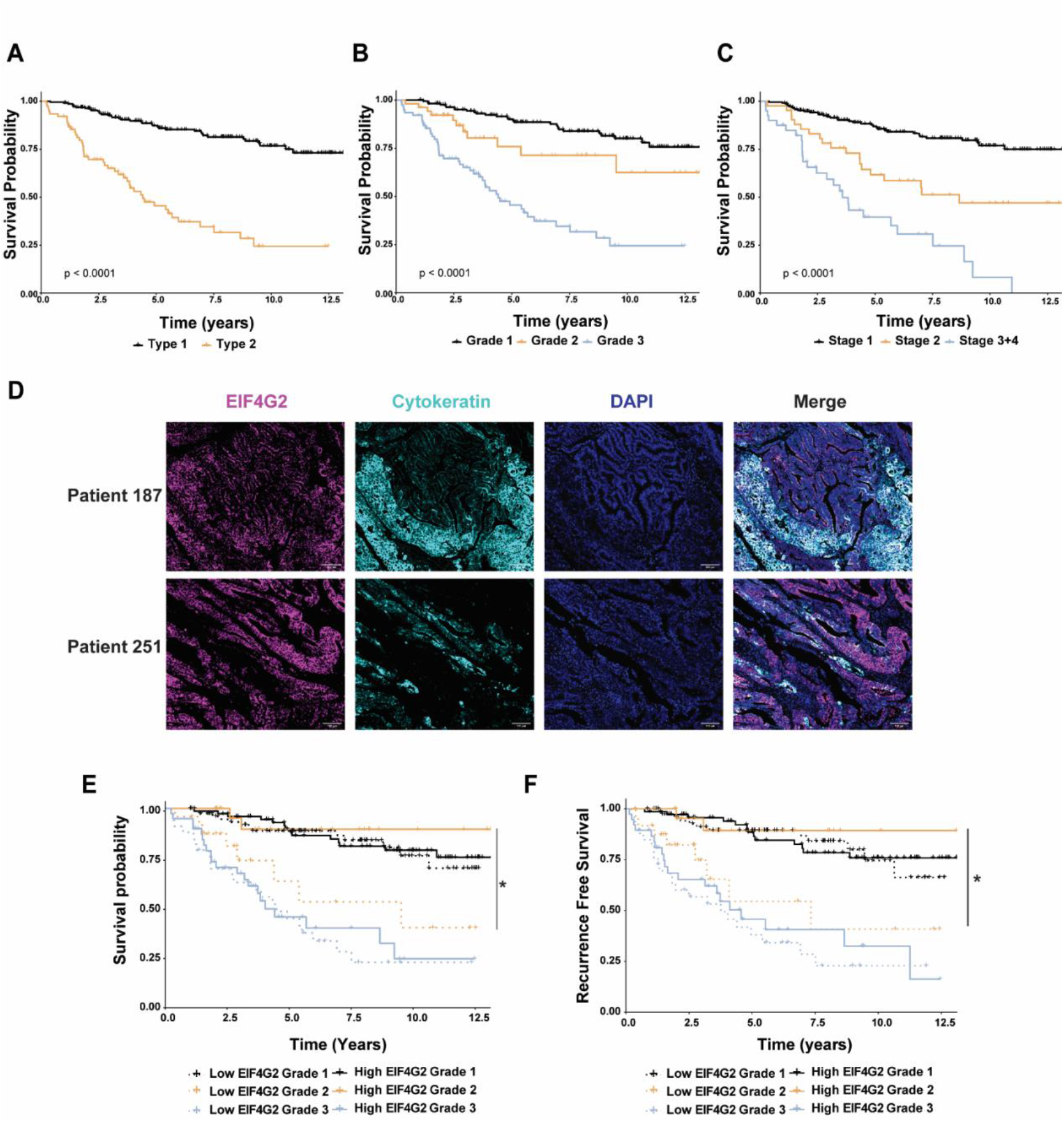
Low EIF4G2 expression correlates with poor prognosis in Grade 2 EC patients. **A-C** Overall survival of 280 endometrial patients according to tumor **(A)** Type, **(B)** stage and **(C)** grade. Kaplan Meier *P*-val was determined using log rank P test. **D** formalin fixed paraffin embedded (FFPE) tumor microarray (TMA) sections from the patient cohort were co-immunostained for EIF4G2, CK and DAPI. Representative images of two patients are shown. Scale bar 100µM. **E, F** EIF4G2 staining within CK positive cells was stratified according to high (above median staining intensity) and low (below median staining intensity) levels and overall survival **(E)** and recurrence free survival **(F)** were assessed by Kaplan-Meier analysis. Log rank *P*-val was calculated from paired comparisons with FDR correction. *: *p*<0.05.

Tissue microarrays (TMA) were generated with cores derived from the 280 re-sectioned formalin-fixed paraffin embedded (FFPE) tumors. Each TMA was immunostained by multiplex immunofluorescence imaging for EIF4G2 and cytokeratin (CK), an epithelial marker used to identify the epithelial tumor cells, and staining intensity was only quantified in CK positive cells. Fig. 1D shows representative staining of EIF4G2 in selected patients. Low and high EIF4G2 expression levels were defined as values below and above the median expression level, respectively (Supplementary Table S1). Expression levels were then correlated with patient survival by Kaplan-Meier survival analysis. Overall, there was no statistically significant difference in OS between patients exhibiting lower or higher intensity of EIF4G2 protein staining, although there was a slight trend towards lower survival in the low expressers (Supplementary Fig. S1A). Similarly, OS and recurrence-free survival did not differ for patients with high or low EIF4G2 expression when analyzed by tumor type (Supplementary Fig. S1B and S1C) or tumor stage (Supplementary Fig. S1D and S1E), although in the latter there was a trend towards decreased OS, especially when comparing EIF4G2 expression in stage 1 and stage 3/4 tumors. However, differences in EIF4G2 expression became statistically meaningful when patients were stratified by tumor grade; there was significantly lower OS in patients with Grade 2 tumors that expressed low levels of EIF4G2, such that the survival curve resembled that of more advanced Grade 3 tumors, whereas the survival curve of high EIF4G2 expressers shifted towards the Grade 1-like curve (Fig. 1E). Following the diagnosis, more than half of the Grade 2 patients were treated with adjuvant therapy in similar numbers, regardless of EIF4G2 expression levels (73% and 68% of the high and low EIF4G2 stratified patients, respectively, Table 1). Treatment was mainly brachytherapy (94.7% of high EIF4G2 and 88.2% of low EIF4G2 stratified patients who received therapy), with a smaller percent of patients receiving chemotherapy, with no statistically significant difference in treatment between the tested groups. The distribution of the Grade 2 tumors according to tumor stage was similar for all expression levels, with the majority in both groups classifying as low stage, meaning the extent of migration and invasion was limited and did not differ between the groups (Table 1). Notably, however, low levels of EIF4G2 expression were associated with increased recurrence rates (24% compared to 15.3% in high EIF4G2), with significantly poorer recurrence-free survival (Fig. 1F). Collectively, these data reveal that lower EIF4G2 expression is associated with poorer outcome in EC based on grade, with an especially strong association between low EIF4G2 expression and poor overall and recurrence-free survival in Grade 2 EC patients.

**Table 1.** Grade 2 patient parameters stratified by low and high EIF4G2 protein Expression.

|  |  | **EIF4G2 Low**<br>**(Intensity<3.7)** | **EIF4G2 High**<br>**(Intensity>3.7)** |
| --- | --- | --- | --- |
| **Patient Data** | **Number of Patients** | 25 | 26 |
|  | **Age** | 65.4 | 62.5 |
|  | **BMI** | 29.0 | 42.6 |
| **Stage** | **1** | 19 (76%) | 19 (73%) |
|  | **2** | 3 (12%) | 5 (19.2%) |
|  | **3** | 3 (12%) | 2 (7.6%) |
|  | **4** | 0 | 0 |
| **Therapies** | **Received Therapy Overall** | 17 (68%) | 19 (73%) |
|  | **Chemotherapy** | 4/17 (23.5%) | 1/19 (5.2%) |
|  | **Brachytherapy** | 15/17 (88.2%) | 18/19 (94.7%) |
|  | **External Radiation** | 7/17 (41.1%) | 5/19 (26.3%) |
| **Survival Statistics** | **Recurrence** | 6 (24%) | 4 (15.3%) |
|  | **Overall Survival (Months)** | 48.6 | 67.1 |
| **Number of Patients from Date of Surgery** | **1 year** | 3 | 0 |
|  | **2-3 years** | 12 | 8 |
|  | **4-5 years** | 1 | 5 |
|  | **6-7 years** | 1 | 2 |
|  | **8-12 years** | 8 | 11 |

### EIF4G2 KD results in increased resistance to anti-cancer agents in HEC-1A and RL95-2 endometrial cell lines

Considering the correlation between Grade 2 EC patient survival and EIF4G2 expression, we wished to determine whether there is a causative link between EIF4G2 and tumor aggression. To this end, EIF4G2 was depleted in the HEC-1A and RL95-2 endometrial carcinoma cell lines. HEC-1A cells were derived from a Stage 1A moderately well differentiated adenocarcinoma (29), and has been classified histologically as Grade 2 (30). RL95-2 cells were derived from Grade 2 moderately differentiated adenosquamous carcinoma (31). Stable knock-down (KD) of EIF4G2 was achieved by viral infection of HEC-1A and RL95-2 cells with vectors expressing shEIF4G2 or shGFP as control (Fig. 2A). Depletion of EIF4G2 slightly reduced the basal growth rates compared to the control cells in both HEC-1A and RL95-2 cells (Fig. 2B). We then tested the effect of EIF4G2 depletion on the responses to Taxol (Paclitaxel), the common chemotherapy agent used against aggressive EC tumors (mostly Type 2 and Grades 2 and 3 tumors) (32). Control and EIF4G2 KD HEC-1A and RL95-2 cells were treated with Taxol for 4 days, and cell viability was measured and compared to DMSO treated cells. While overall, RL95-2 cells were more responsive to the treatment than HEC-1A cells, in both cell types EIF4G2 KD cells exhibited increased viability in the presence of Taxol compared to the control KD cells (Fig. 2C). Control and EIF4G2 KD HEC-1A and RL95-2 cells were also exposed to X-ray irradiation, a front-line therapy given either internally (brachytherapy) or externally. A lower dosage was used for the RL95-2 cells since they proved more sensitive to irradiation. EIF4G2 depletion increased the viability in irradiated HEC-1A (16 Gy) and RL95-2 (8 Gy) cells by ∼20% and ∼50%, respectively, compared to the control KD cells (Fig. 2D). To assess the molecular effects of irradiation on the cell populations, western blotting was performed for γH2AX, a histone variant that is phosphorylated in response to double stranded DNA breaks and serves as a marker for DNA damage and resolution. In both cell lines, EIF4G2 KD cells showed reduced levels of γH2AX compared to the control cells (Fig. 2E and 2F, Supplementary Fig. S3E). This suggests that the irradiated EIF4G2 KD cells either underwent less DNA damage or managed to resolve the damage faster and/or more efficiently compared to control irradiated cells. Overall, EC cells that are depleted of EIF4G2 displayed reduced sensitivity to standard therapies used clinically to treat EC.

**Fig. 2.**
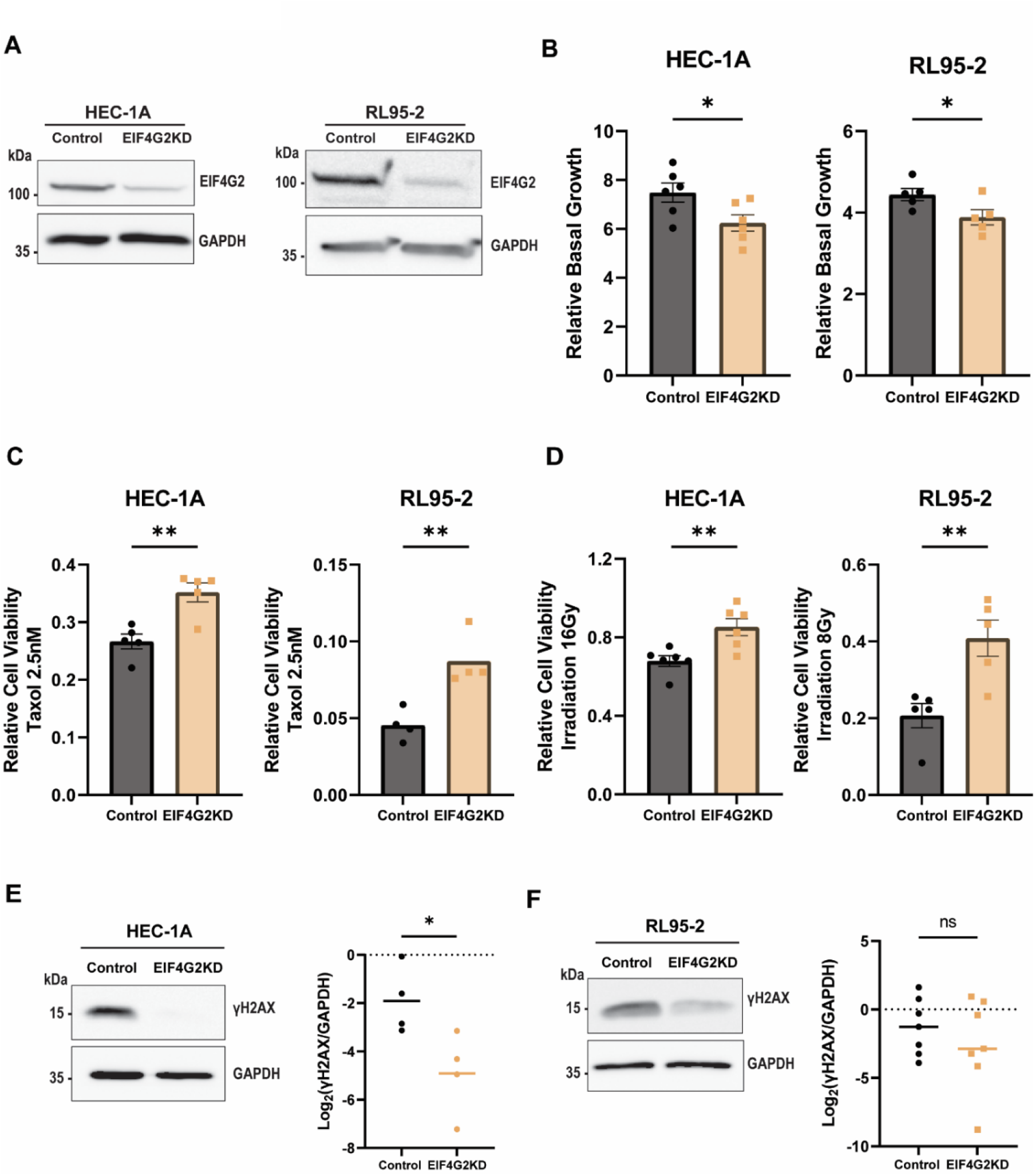
Depletion of EIF4G2 in HEC-1A and RL95-2 cells results in increased therapy resistance. **A** Total cell lysate form HEC-1A (left) and RL95-2 (right) control and EIF4G2 KD cells were subjected to western blot analysis for EIF4G2 and GAPDH as loading control. Representative blots are shown. **B** Relative basal growth of control and EIF4G2 KD cells was measured by CellTiter-Glo assay in HEC-1A (left) and RL95-2 (right). Relative growth was calculated as luminescence of cells at day 4 (T4) relative to corresponding cells at day 0 (T0). Shown are the mean relative luminescence values±SEM of n>5 independent experiments. Significance was determined by two tailed t-test. *: *p*<0.05. **C** Cell viability was measured by CellTiter-Glo assay in HEC-1A (left) and RL95-2 (right) control and EIF4G2 KD cells incubated with DMSO or 2.5 nM Taxol for 4 d. Luminescence values of Taxol treated samples were normalized to the values of corresponding DMSO treated samples, representing relative viability. Shown are the mean relative luminescence values±SEM of n>4 independent experiments. Significance was determined by two tailed t-test. **: *p*<0.01. **D** HEC-1A (left) and RL95-2 (right) control and EIF4G2KD cells were exposed to either 16G or 8G X-ray irradiation, respectively. Cell viability was assayed 4 d later as in (**C**) relative to mock irradiated cells. Shown are the mean relative luminescence values±SEM of n>5 independent experiments. Significance was determined by two tailed t-test. **: p<0.01. **E, F** Total cell lysates form HEC-1A **(E)** and RL95-2 **(F)** control and EIF4G2 KD cells were subjected to western blot analysis for γH2AX and GAPDH, as loading control, 4 d following irradiation. γH2AX signal was normalized to GAPDH and quantification results are represented as individual data points and also as mean values of 4 independent experiments **(E)** or 7 independent experiments **(F)**.

### EIF4G2 depletion increases the proportion and treatment resistance of cells expressing markers of aggression CD133 and CD44

In order to further characterize the effect of EIF4G2 protein depletion on cancer outcome, and to understand why EIF4G2 KD cells are more resistant to therapies, we examined the prevalence of aggressive sub-populations within the control and EIF4G2 KD cells. The most prominent markers for identification and isolation of these aggressive populations from both primary tumors and cell lines are CD133 (PROM1), CD44 and ALDH1A1^high^ expression. The effect of EIF4G2 depletion on CD133 and CD44 cell surface expression was assessed by FACS analysis (CD133 and CD44 gating strategy shown in Supplementary Fig. S2A and 2B, respectively). While the vast majority (>96%) of HEC-1A cells did not express CD133, significantly, EIF4G2 KD increased both the marker’s expression intensity and the population distribution of CD133 positive cells, nearly doubling from 3.4% to 5.9% positive cells (Fig. 3A). CD44 was expressed at varying levels in most cells, and again, EIF4G2 KD resulted in a shift towards higher expression, enhancing both the percent of cells expressing high levels of the marker, and the intensity of the expression (Fig. 3B). Similar, yet more modest, results were also observed upon transient siRNA transfection (Supplementary Fig. S3A and S3B). RL95-2 cells did not express CD133, even upon EIF4G2 KD, yet EIF4G2 depletion did increase CD44 surface expression intensity by two-fold (Fig. 3C), similar to the HEC-1A cells.

**Fig. 3.**
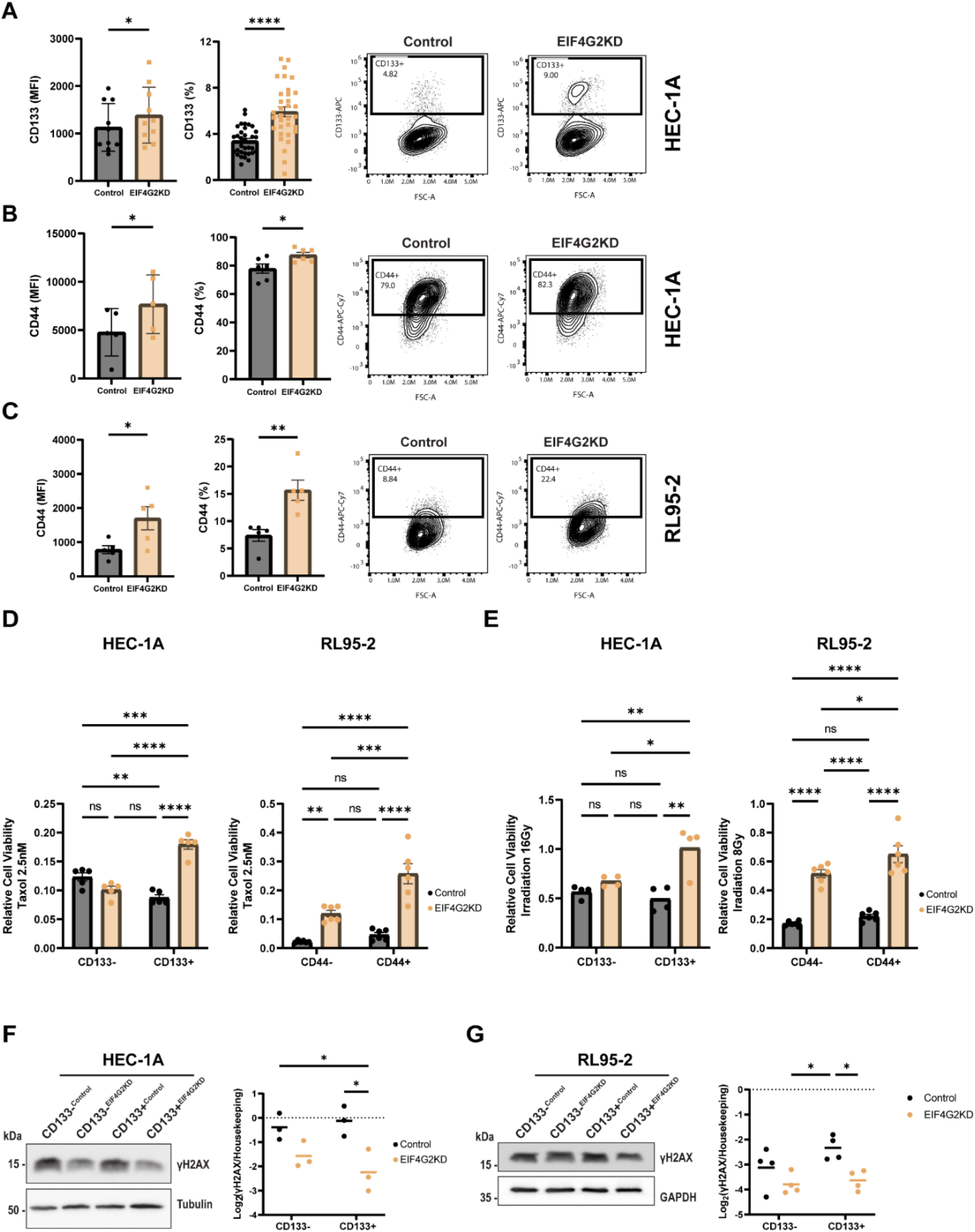
EIF4G2 KD increases CD133 and CD144 expression and viability of CD133+ and CD44+ populations following therapy in HEC-1A and RL95-2 cells. **A-C** Flow cytometry analysis of cell-surface expression of CD133 **(A)** and CD44 **(B, C)** in control and EIF4G2 KD HEC-1A cells (**A, B**) and control and EIF4G2 KD RL95-2 cells (**C**). Quantification of results is presented as mean±SEM, n>5 for both overall mean florescent intensity (MFI) and as percent of total population. *P*-vals were calculated using two-tailed t-test (*: *p*<0.05, **: *p*<0.01, ****: *p*<0.0001). A representative flow cytometry plot is shown for each. **D** Control and EIF4G2 KD HEC-1A and RL95-2 cells were separated for CD133 and CD44 expression, respectively. Separated populations were treated with 2.5 nM Taxol or DMSO as control for 4 d and cell viability assayed by CellTiter-Glo. Luminescence was normalized to DMSO treated cells. Left graph, HEC-1A cells, right graph RL95-2 cells. **E** Cell viability of HEC-1A (left) and RL95-2 (right) CD133+ and CD133-control and EIF4G2 KD separated populations 4 d after exposure to 16Gy (HEC-1A) or 8Gy (RL95-2) X-ray irradiation. For (**D, E**), results are presented as mean values±SEM of n>4 independent experiments. Significance was determined by two-way ANOVA. (*: *p*<0.05, **: *p*<0.01, ***: *p*<0.001, ****: *p*<0.0001). **F, G** Total cell lysates form HEC-1A **(F)** and RL95-2 **(G)** CD133+ and CD133-control and EIF4G2 KD separated populations were subjected to western blot analysis for γH2AX and GAPDH, as loading control 4 d following irradiation. γH2AX signal was normalized to GAPDH and quantification results are presented as individual data points and also as mean values of 4 independent experiments. Statistical significance was determined for log_2_ transformed mean values by two-way ANOVA (*: *p*<0.05).

Since a more pronounced resolution of the highly expressing cell populations was observed with the CD133 marker in HEC-1A cells, the potentially more aggressive sub-populations were separated from the KD cells based on this marker using anti-CD133 conjugated magnetic beads. Generally, there was no significant difference in the proliferation rate of the separated populations (Supplementary Fig. S3C). Culturing over time following separations did not significantly alter the percentage of the CD133^+^ or CD133^-^ populations (Supplementary Fig. S3D). Likewise, the RL95-2 cells were separated according to the CD44 marker, using anti-CD44 conjugated magnetic beads.

Cell viability of CD133+ separated HEC-1A and CD44+ separated RL95-2 control and EIF4G2 KD cells was assessed following treatments with Taxol and X-ray irradiation. CD133+^EIF4G2KD^ cells displayed a significant increase in viability compared to all other separated populations in response to Taxol treatment. Similar results were observed with the CD44+ RL95-2 cells (Fig. 3D). A pronounced effect was also detected after irradiation with X-rays. The relative number of viable cells in the CD133+ EIF4G2 KD cells was almost double that of the other cell populations in HEC-1A cells, and almost three times more than the control populations in the RL95-2 cells (Fig. 3E). The cell lysates of irradiated cells 4 d after treatment were blotted for γH2AX. As expected, γH2AX was activated, yet remained significantly lower in EIF4G2KD CD133+ HEC-1A and CD44+ RL95-2 cells (Fig. 3F and 3G). This implies that DNA damage was reduced and/or more quickly resolved in the EIF4G2 KD cells, and thus these more aggressive sub-populations were more likely to survive the irradiation.

### Downregulation of EIF4G2 expression alters the transcriptomic and proteomic signature of CD133+ cells and selectively impairs translation of kinesin 1 proteins

In order to better understand the phenotypic differences in response to therapies, in particular the increased resistance of the CD133+ cell population, control and KD HEC-1A populations were FACS sorted and then subjected to RNA-seq transcriptomic analysis and mass spectrometry (MS) analysis. The CD133- and CD133+ control populations showed relatively minor changes in their transcriptomic profiles with only 77 differentially expressed genes (DEGs) (Supplementary Table S2). Annotation by GeneAnalytics (33) of this set indicated enrichment in pathways associated with “cytoskeletal signaling” (7 genes) and “embryonic and induced pluripotent stem cell and lineage specific markers” (5 genes: SOX17, TNNI3, PAX2, PROM1 (CD133), FGF18) (Supplementary Fig. S4A), consistent with a more stem-like phenotype in the CD133+ cells. There was also a high degree of similarity between the transcriptomic profiles of the CD133-populations regardless of EIF4G2 status (137 DEGs, with no high scoring pathways by GeneAnalytics), indicating little effect of EIF4G2 depletion on the general HEC-1A cells. In contrast, the RNA-seq analysis highlighted a strong transcriptomic difference between the CD133+^EIF4G2KD^ cells and all other populations (Fig. 4A). In other words, EIF4G2 depletion had the largest effects on gene expression in the context of the CD133+ cells, with 939 DEGs between the EIF4G2 and control CD133+ populations. Moreover, its depletion amplified the relatively small transcriptomic differences between the CD133- and CD133+ cells by more than 10-fold (77 DEGs vs 894 DEGs in the EIF4G2KD comparison) (Supplementary Table S2). Gene annotation of both sets of DEGs for significant pathways with high score revealed pathways related to cellular metabolism. In addition, the malignant pleural mesothelioma pathway and the glucocorticoid receptor pathway showed enrichment in the CD133+^EIF4G2KD^/CD133+^Control^ and CD133+^EIF4G2KD^/CD133-^EIF4G2KD^ comparisons, respectively (Supplementary Fig. S4B and S4C). Notably, upon EIF4G2 KD, ALDH1A1, a prominent marker whose up-regulation is associated with enhanced tumor aggressiveness and therapy resistance in various cancers, including EC (34–38), was increased in expression in the CD133+ cells. Thus decreased EIF4G2 expression not only enhanced the prevalence of the aggressive populations (in terms of CD133 and CD44 expression), but also changed their transcriptomic profile compared to CD133+ cells.

**Fig. 4.**
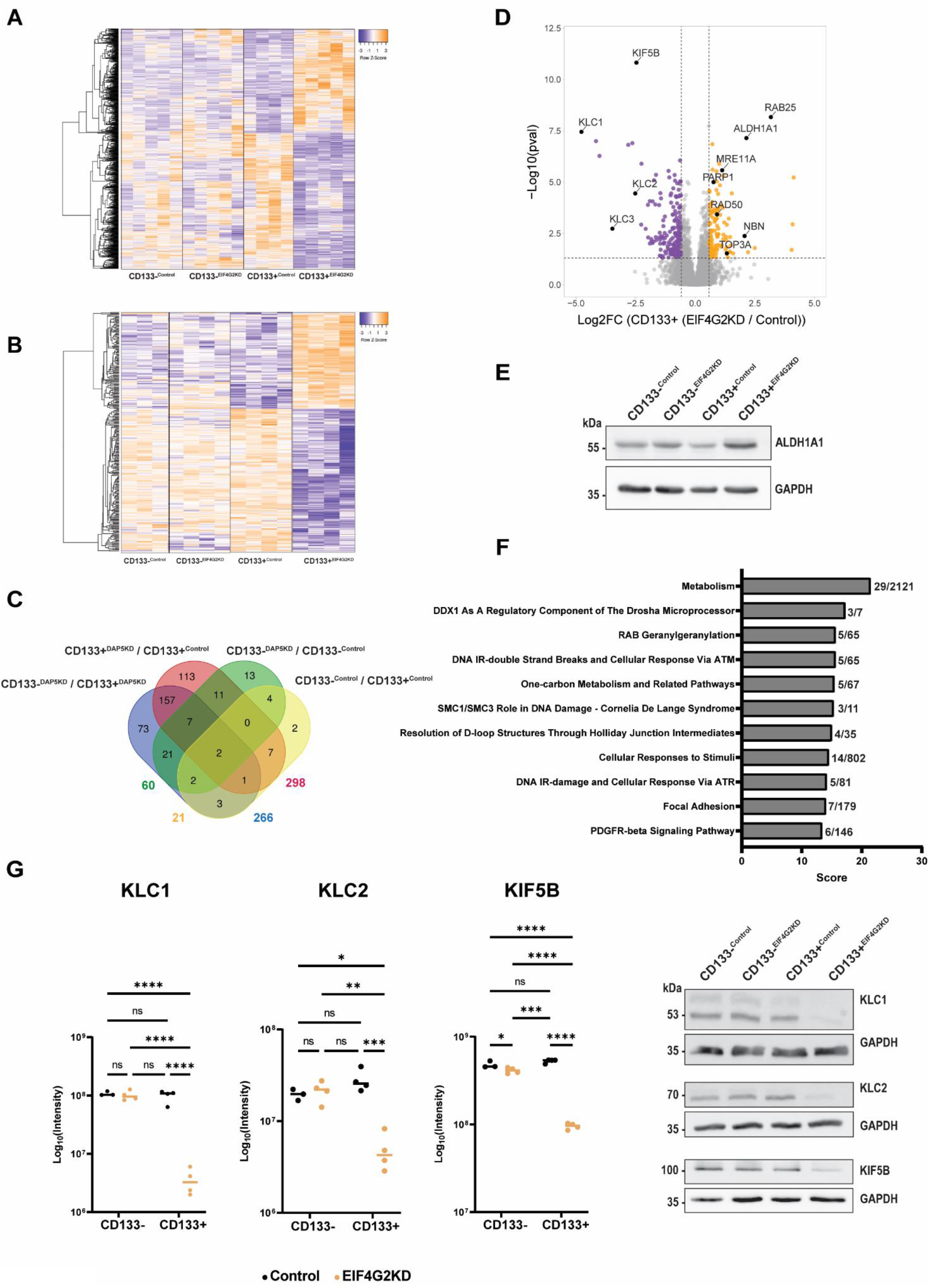
EIF4G2 KD alters the transcriptomic and proteomic signatures of CD133+ population in HEC-1A cells. Control and EIF4G2 KD HEC-1A cells were FACS sorted by CD133 marker expression and subjected to RNA-seq and MS analysis. **A** Heat map showing hierarchical clustering of gene expression levels of all DEGs identified following RNA-seq analysis of CD133- and CD133+ sorted control and EIF4G2 KD populations. **B** Heat map showing hierarchical clustering of all proteins with significantly changed abundance in MS analysis of CD133- and CD133+ sorted control and EIF4G2 KD HEC-1A cells. **C** Venn diagram showing overlap of all differentially abundant proteins among the four comparisons. Numbers at the edges of the diagram represent the overall number of proteins with differential abundance in each comparison. **D** Volcano plot of the Log_2_ (Fold Ratio) of the abundance of the detected proteins in CD133+^EIF4G2KD^ / CD133+^Control^ comparison, vs. their significance expressed as log_10_ *p*-value. Proteins with significant increased abundance are indicated in orange, and decreased abundance in purple. **E** Total cell lysates from separated control and EIF4G2 KD HEC-1A populations were subjected to western blot analysis for ALDH1A1 and GAPDH as loading control. Shown is a representative blot of 3 independent experiments. **F** High scoring significant pathways identified by GeneAnalytics pathway analysis of the set of proteins with increased abundance in the CD133+^EIF4G2KD^ / CD133+^Control^ comparison. Score numbers on X-axis indicate significance, numbers at right represent the number of proteins identified in the dataset out of the total number of proteins within the given pathway. **G** Protein expression of KLC1, KLC2 and KIF5 based on the abundance detected by the MS analysis. Data presented as mean of Log_10_ (Intensity), n=4. Statistical significance was determined by two-way ANOVA (*: *p*<0.05, ***: *p*<0.001, ****: *p*<0.0001). Representative western blots of HEC-1A separated CD133- and CD133+ control and EIF4G2 KD cells validating the MS results for KLC1, KIF5B, and KLC2. GAPDH was used as a loading control. Shown is one representative blot of n=3.

The phenotypic and transcriptomic effects of EIF4G2 KD on CD133+ cells are likely ascribed to its role as a translation factor, as has been shown in ESCs (15). Polysome profiling analysis of EIF4G2 and control KD HEC-1A cells showed no changes in global translation (Supplementary Fig. S5A), consistent with EIF4G2’s involvement in selective translation of specific mRNA targets (9,14,15,21). Furthermore, investigation of the proteome of EIF4G2 KD CD133- and + cells by MS confirmed that the majority of proteins were not changed in abundance. An overall comparison of the proteomic signatures of the CD133- and + cells with or without EIF4G2 KD resembled the transcriptomic pattern, in that the CD133+^EIF4G2KD^ proteome was altered compared to the other 3 sorted populations, which all shared a similar profile (Fig. 4B). Enriching for CD133 did not in itself result in major differences in the proteome; only 21 proteins changed in abundance in the control CD133+ vs CD133-populations (Fig. 4C). The difference between the populations was enhanced by EIF4G2 KD, with 266 proteins exhibiting changed abundance in the CD133+^EIF4G2KD^ vs CD133-^EIF4G2KD^ cells. EIF4G2 KD had a moderate effect on the proteome of CD133-cells, with 60 differentially abundant proteins in the KD vs control cells, and a stronger effect on the CD133+ cells, with 298 proteins with differential abundance identified when comparing CD133+^EIF4G2KD^ and CD133+^Control^ populations (Fig. 4C). Thus, EIF4G2 KD affected both the transcriptomic and proteomic profiles of the CD133+ cells to the greatest extent.

Of the 298 proteins with differential abundance in the CD133+^EIF4G2KD^ and CD133+^Control^ comparison, 20 were also observed to change in the CD133-^EIF4G2KD^ and CD133-^Control^ comparison, indicating a dependence on EIF4G2 expression unrelated to the cell population. The majority of changes, however, occurred due to both the CD133 and the EIF4G2 KD status. Of the 298 differentially abundant proteins in the CD133+ control and KD cells, 177 showed decreased abundance and 121 were increased. Pathway analysis of the decreased set of proteins by GeneAnalytics showed enrichment in pathways associated with the mitochondrial electron transport chain, metabolism, cell adhesion, actin nucleation and G-protein/Rho GTPase signaling pathways (Supplementary Fig. S5B).

The MS analysis showed a set of 121 proteins with increased abundance specifically in the EIF4G2 KD CD133+ cells compared to the control CD133-cells (Fig. 4D). While these do not represent potential EIF4G2 mRNA translation targets, they do shed light on the indirect contribution of EIF4G2 to endometrial cancer aggressiveness, particularly within CD133+ cells. As observed for its mRNA, the tumor aggressiveness marker ALDH1A1 increased more than 4-fold in the MS analysis (Fig. 4D). This increase was validated by western blotting (Fig. 4E). No difference was observed between control CD133- and + cells in either the MS dataset or the western blot. GeneAnalytics pathway analysis of the 121 proteins with increased abundance showed they were enriched in the general pathway “metabolism” and several related pathways with the same 3-5 proteins involved in DNA repair of double strand breaks and response via ATM (Fig. 4F). These included PARP1, RAD50, MRE11A, TOP3A and NBN (also known as NBS1) (Fig. 4D).

Of the 177 proteins with decreased abundance, 126 did not show comparable decreases in mRNA expression (Supplementary Fig. S5C), indicating that the decreased protein levels observed were not due to changes in gene transcription or mRNA stability. While not excluding other factors that can influence protein steady state levels, this set of proteins may represent potential translation targets of EIF4G2. In fact, several of these proteins have been previously identified as EIF4G2 translation targets by unbiased screens using either ribosome footprinting (9,14,21) or polysome profiling (15) in additional cell types. These included KIF5B (9), which was among the highest scoring proteins with changed abundance in the EIF4G2 KD CD133+ cells. KIF5B is a kinesin heavy chain, which either alone or together with a pair of light chains, forms the kinesin 1 microtubule-based motor protein. Notably, 2 of these light chains, KLC1 and KLC2, also decreased in abundance upon EIF4G2 KD in the CD133+ cells (Fig. 4D), and a third light chain, KLC3, showed reduced abundance upon EIF4G2 KD in both CD133- and CD133+ cells. The MS results were confirmed by western blot analysis of these proteins; KLC1, KLC2 and KIF5B steady state levels strongly decreased in the CD133+^EIF4G2KD^ population compared to all others (Fig. 4G). Thus, EIF4G2 is necessary to maintain the steady state levels of kinesin 1 in HEC-1A CD133+ cells.

### Low protein staining intensity of EIF4G2 dependent targets KLC1 and KIF5B correlates with poor patient survival

Considering the prominent decrease in kinesin 1 motor proteins upon EIF4G2 KD, we wished to extend this observation and assess their status in EC patients. Sequential sections from the TMAs of the patient cohort used in Fig. 1 were stained for KLC1 and KIF5B together with CK by multiplex immunofluorescence. Fig. 5A shows representative staining from representative patients. Expression levels based on quantification of staining intensity were compared to EIF4G2 staining of the next section from the same TMA cores. Notably, there was a small but significantly positive correlation between expression of EIF4G2 and KLC1, and a moderately larger positive correlation with KIF5B expression (Supplementary Fig. S6A and S6B). OS was then correlated with expression of KLC1 and KIF5B, with low and high expression defined as values that were less than or greater than the median staining intensity for each protein, respectively. In general, OS curves trended lower in patients with low KLC1 or low KIF5B expressing tumors upon stratification of tumors by type, grade and stage, although this trend was not always statistically significant (Fig. 5B-G). Notably, OS was significantly reduced in Type 1 and Grade 2 tumors with low KIF5B staining intensity (Fig. 5C, G), and in low KLC1 expressing tumors of advanced Stages 3/4 (Fig. 5D). Notably, the shift in patient survival probability in Grade 2 tumors expressing low KIF5B (and also KLC1, although this was not statistically significant) towards the Grade 3 curve was reminiscent of the OS curve of Grade 2 low expressing EIF4G2 patients (Fig. 5F, G, Fig. 1E). Recurrence-free survival analysis (Supplementary Fig. S6C-H) revealed a significant decrease in Grade 2 patients with low KLC1 expression (Supplementary Fig. S6G), and in Type 1 and Grade 2 patients with low KIF5B staining intensity (Supplementary Fig. S6D, H). Overall, these results show a strong correlation between low protein expression of either KLC1 or KIF5B and poorer survival outcomes in specific EC tumor classes.

**Fig. 5.**
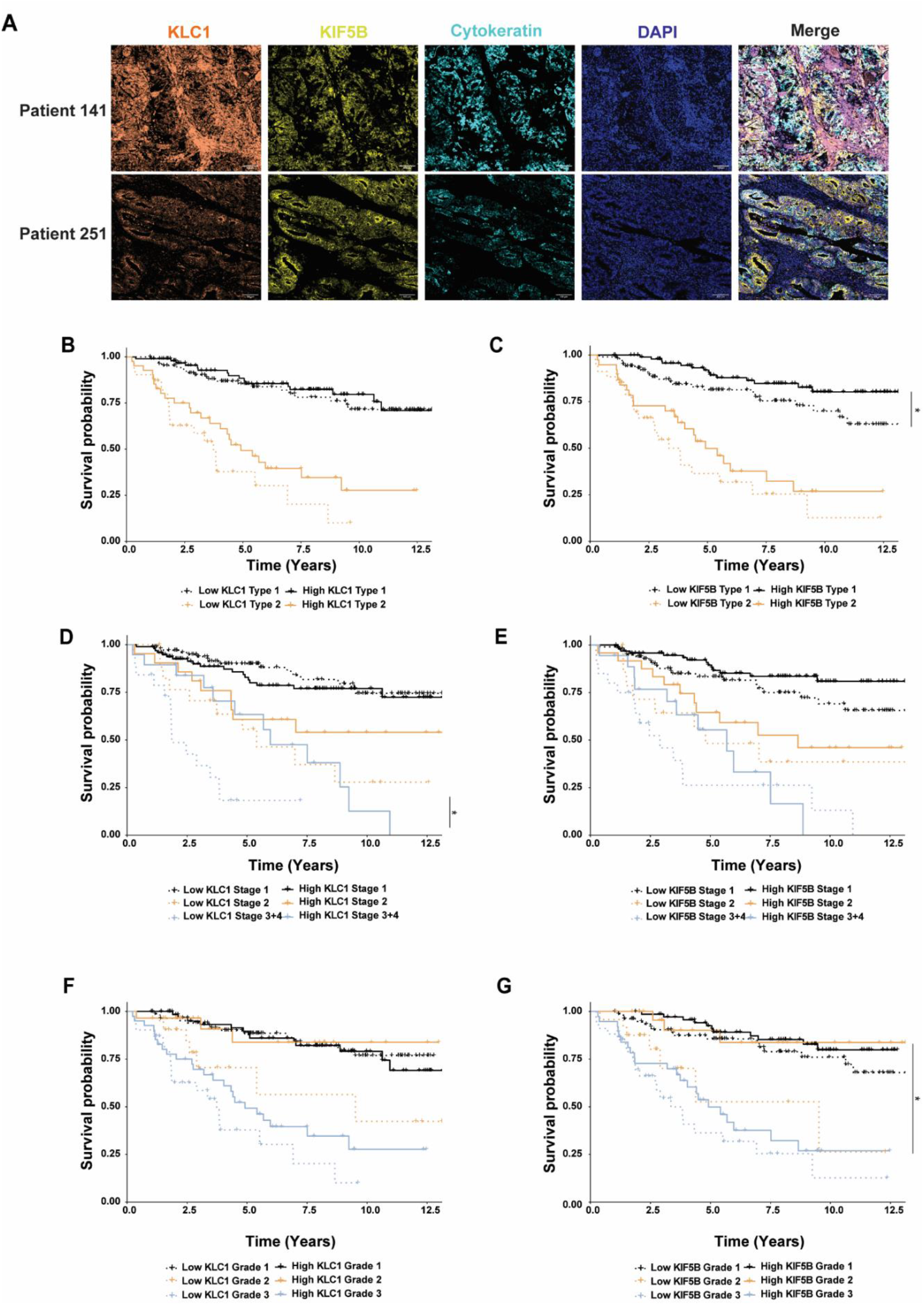
Low protein expression of EIF4G2 targets KLC1 and KIF5B correlates with worse overall survival in EC patients. **A** FFPE tumor TMA sections from the 280 patients were immunostained for KLC1, KIF5B, CK and DAPI. Representative images of two patients are shown. Scale bar 100µM. **B-G** Overall survival of the 280 endometrial patients according to tumor **(B, C)** type, **(D, E)** stage, and **(F, G)** grade. KLC1 **(B, D, F)** and KIF5B **(C, E, G)** staining in CK positive cells was stratified according to high and low intensity levels compared to the calculated median, and overall survival was assessed by Kaplan-Meier statistics. Log rank *P*-val was determined for protein expression of each of these signatures. Paired comparison was calculated with FDR correction. *: *p*<0.05, ***: *p*<0.001.

## Discussion

Here we examined the contribution of the non-canonical translation factor EIF4G2 to endometrial cancer in human patients and cell lines. EC cells depleted of EIF4G2 were less sensitive to Taxol and radiation, common therapeutic agents used clinically to treat EC. Furthermore, EIF4G2 KD resulted in enrichment in cells expressing higher levels of markers such as CD133, CD44 and ALDH1A1 associated with aggressiveness, often referred to as cancer stem cells (CSC). This sorted cell population had a vastly changed transcriptome and proteome upon EIF4G2 depletion, indicating an inherently different cell phenotype, and was particularly resistant to both chemotherapy and radiotherapy. The resistance to therapy is consistent with the increased ALDH1A1 expression at both the mRNA and protein levels. Specifically, ALDH1A1 overexpression was shown to be necessary for Paclitaxel resistance in EC cells grown as spheroids *in vitro* (35). The resistance to radiation may be connected to increased expression of several DNA damage repair proteins, including PARP1, and components of the MRN (MRE11, RAD50, and NBN) complex that senses and repairs double stranded DNA breaks. In fact, lower levels of γH2AX in EIF4G2 KD cells are consistent with an improved ability to resolve DNA damage. Thus, KD cells are better equipped to deal with DNA damage and are more likely to survive DNA damaging treatments. This infers a survival advantage to tumor cells that display lower expression levels of EIF4G2 expression, with implications for enhanced drug resistance and/or recurrence.

Overall, the data predict that loss of or decreased EIF4G2 expression, which has been previously documented in EC (BioRxiv 2023.08.22.554280), would significantly lead to poorer clinical outcome. Notably, our analysis of EIF4G2 protein levels in the EC tumors indicated that low expression correlated with poorer outcome and increased recurrence rates specifically in patients with Grade 2 tumors. While the clinical data did not clarify as to why EIF4G2 levels would be critical specifically for this tumor grade, it is consistent with EIF4G2’s previously reported divergent association with different types and stages of cancers ((22–24) BioRxiv 2023.08.22.554280). Importantly, Grade 2 EC tumors are usually considered low risk and are not treated aggressively, yet some manifest clinical indications more reminiscent of higher Grade 3 tumors and should be treated as such. EIF4G2 appears to be a promising prognostic marker to differentiate between non-aggressive and more aggressive Grade 2 tumors. Based on our data, Grade 2 tumors with low EIF4G2 levels would benefit from more aggressive adjuvant treatment. In fact, while most of the Grade 2 patients in our cohort were treated with radiation therapy, only a small number received chemotherapy. Considering the relative resistance of EIF4G2 depleted EC cells to Taxol and radiation *in vitro*, even when these treatments were applied, they may have been limited in their efficacy in tumors with low EIF4G2 expression. Further assessment of the drug response to these different treatments in tumors with varying levels of EIF4G2 is mandated to clarify the clinical implications of our current findings.

Our findings on the tumor suppressive nature of EIF4G2 in EC contradict previous in-depth analysis of EIF4G2 function in metastatic triple negative breast cancer, in which EIF4G2 was shown to promote metastasis through translation of factors associated with EMT, motility, angiogenesis and cell survival (22). Moreover, EIF4G2 protein levels were elevated in metastatic tumors compared to non-metastatic tumors from a very small cohort of triple negative breast cancer patients, and high mRNA levels correlated with poor metastasis-free survival rates (22). Our much wider protein analysis of a larger group of primary tumors surgically extracted from EC patients did not reveal adverse survival probabilities associated with increased EIF4G2 protein expression, but rather the opposite. In addition to the fact that immunostaining for EIF4G2 protein in tissue is a more accurate description of the relative levels of EIF4G2 specifically within the tumor tissue compared to measurement of total mRNA levels, the discrepancy in the data is consistent with EIF4G2 having different functions in different cancer subtypes. In breast cancer too, EIF4G2 depletion affected metastasis but not growth of primary tumors upon injection of EIF4G2 KD cells into mice (22). Thus EIF4G2’s role, and by extension its direct translation targets, appears to be context dependent, with divergent roles in metastatic and primary tumors.

While our study did not directly determine mRNA translation targets of EIF4G2, our combined transcriptomic and proteomic analysis of EIF4G2 KD CD133+ EC cells does provide clues to potential targets among proteins with decreased abundance but unchanged mRNA levels. Downregulated pathways in the MS analysis included pathways related to adhesion, cytoskeleton and integrins, in line with some of the translation targets identified in breast cancer. Other potentially relevant pathways with changes were related to the electron transport chain of oxidative respiration and energy metabolism. Four NADH dehydrogenase (complex I) proteins, one component of succinate dehydrogenase (complex II), two ubiquinol-cytochrome c reductase (complex III) proteins and COX16, which is necessary for assembly of cytochrome oxidase (Complex IV), as well as additional proteins involved in mitochondrial protein import or morphology, were all reduced in abundance in EIF4G2 KD CD133+ cells (see Supplementary Table S3). These results were reminiscent of hESCs, in which mitochondrial oxidative phosphorylation proteins exhibited reduced translation upon EIF4G2 KD, accompanied by a defective oxidative respiratory pathway (15). Pluripotent stem cells are usually more glycolytic, and differentiation is accompanied by increased reliance on mitochondrial oxidative respiration. Thus, impaired oxidative respiration can limit the transition from pluripotency to more differentiated states (39).

In general, cancer cells are known to rely heavily on glycolysis for energy production (referred to as the Warburg effect), and in some cancers, such as lung, colon, breast and ovarian cancers, osteosarcoma and glioblastoma, CSCs are even more glycolytic compared to their more differentiated counterparts (40). Endometrial cancer spheroid cells with CSC properties also showed enhanced glycolysis that was dependent on high ALDH activity and ALDH1A1 expression (35). It therefore follows that the predicted downregulation of the oxidative phosphorylation pathway in EIF4G2 KD EC cells likely confers a more stem-like phenotype to these cells. This, combined with increased expression of markers CD133, CD44 and ALDH1A, and the enhanced therapy resistance of these cells, suggests that similar to its role in ESCs, EIF4G2 is necessary for the dynamic transition between more aggressive stem-like and more differentiated EC cells.

As a direct consequence of EIF4G2 depletion in the CD133+ EC cells, decreased protein abundance was observed for components of the Kinesin 1 microtubule motor protein, including heavy chain KIF5B and light chains KLC1, 2, 3. One of these, KIF5B, has been shown to be a direct translation target of EIF4G2 in two different cell systems by two different translation assays (9,22). Interestingly, various mitotic kinesin motor proteins are up-regulated in tumors, and as they play an important role in microtubule dynamics during mitosis and cytokinesis, are targets of anti-cancer drug therapy (41). KIF5B, however, is a non-mitotic member of the kinesin protein superfamily, and either alone or together with its associated light chains, transports organelles such as mitochondria and endosomes towards the plus end of the microtubules. It has also been shown to be involved in microtubule-dependent mobility of DNA with double strand breaks and positioning of the nuclear envelope to facilitate DNA repair (42) (BioRxivs 2023.05.07.539750). Interestingly, similar to EIF4G2, KIF5B seems to have divergent roles depending on the tumor context; in triple negative metastatic breast cancer cells, KIF5B levels were increased, leading to acquisition of EMT, invasiveness and stem-like phenotype (43,44), but silencing of KIF5B in epithelial MDCK cells induced EMT, and enhanced invasive and tumorigenic properties (45). In the former system, KLC1 levels inversely correlated with KIF5B, and was localized to the cytoplasm while KIF5B localized to the nucleus (43), implying that KIF5B partners with other KLCs or functions independently of KLCs in this system within the nucleus to mediate its pro-tumorigenic effects. In our EC TMA samples, in contrast, both KIF5B and KLC1 localized to the cytosol, and their expression positively correlated. Moreover, low expression levels of both proteins were associated with poor OS and recurrent free survival outcomes in the EC patients of specific type, grade and stage. Thus, the kinesin 1 motor protein is likely to play a different role in EC than it plays in metastatic breast cancer, and its expression may serve as a novel prognostic marker in EC, similar to EIF4G2 itself. Our functional analysis of EC cells *in vitro* coupled with expression analysis of a large cohort of EC patients underscores the potential significance of the EIF4G2-kinesin-1 axis for the development of EC. Furthermore, our findings strongly support the integration of these markers into clinical practice for the management of endometrial carcinoma. This strategic inclusion will empower healthcare professionals to tailor prognosis and treatment plans to each patient’s unique profile, ultimately enhancing patient outcomes and the overall quality of care.

## Materials and Methods

### Ethics statement

This study was performed in line with the principles of the Declaration of Helsinki. Patients’ samples and data were collected following approval by the Emek Medical Center Institutional Review Board (IRB, protocol no 0043-22-EMC).

### Human patient samples and tissue microarray construction

Tissue microarrays (TMA)-containing cores from 280 endometrial patients, were retrieved from the archives of HaEmek Medical Center under IRB, protocol no 0043-22-EMC. All clinical data were collected following appropriate ethical approvals. We excluded from our cohort patients with synchronous tumors and patients who received therapies prior to surgery. All patients were females with dates of birth, height and weight stated in Supplementary Table S1. No randomization or blinding was necessary for patient selection. Staining and analysis of patient samples was blinded. Tissue cores were obtained from the original formalin fixed paraffin embedded (FFPE) tissue blocks and from tissue stocks kept in the Pathology Department of Emek Medical Center. All tissue blocks were constructed by using a TMA grand master system (3DHistech Ltd., Budapest, Hungary). Each case underwent precise evaluation by a pathologist to identify the tumor from which the core could be extracted. Up to 60 2 mm cores were placed in each TMA block for a total of six TMA blocks. The first 4-μm section from each block was used for hematoxylin and eosin staining.

### Cell lines and cell culture

HEC-1A (HTB-112) and RL95-2 (CRL-1671) were obtained from the ATCC. Cells were cultured in DMEM/F-12 (Gibco 21331-020) medium with 10% FBS (Gibco 12657-029), 1% penicillin streptomycin (Biological Industries 03031-1B) and 1% L-Glutamine (Biological Industries 03-020-1B). Stable KD of EIF4G2 was generated by infecting HEC-1A or RL95-2 cells with lentiviruses harboring pLKO.1-puro plasmid expressing shRNA targeting GFP (Control) or shRNA targeting EIF4G2 (Sigma TRCN0000147914), followed by selection using puromycin, as previously described (15). Transient EIF4G2 KD was generated by transfecting HEC-1A cells with control siRNA (Dharmacon D-001810-10) or siRNA targeting EIF4G2 (Dharmacon L-011263-00-0020) using JetPEI transfection reagent (Polyplus 101000020) according to manufacturer’s protocol.

### Flow cytometry analysis and sorting

For FACS analysis, 0.5-2 million cells were trypsinyzed, collected and washed. Cells were stained with anti CD133-APC conjugated antibody ((Thermo Fisher Scientific Cat# 17-1338-42, RRID:AB_1603199), 1:100) or CD44-Alexa Fluor 700 ((Thermo Fisher Scientific Cat# 56-0441-82, RRID:AB_494011), 1:400) for 30 min at 4°C. Cells were counterstained with propidium iodide (PI, Sigma-Aldrich, P4864) to assess viability. Data acquisition was performed on CytoFLEX flow cytometer and software (Beckman Coulter Life Sciences) and data analysis was performed using FlowJo software version 10.8.1.

Cell sorting was performed on Control and EIF4G2KD HEC-1A cells following magnetic beads separation of CD133+ and CD133-(CD133 Microbead Kit, Miltenyi Biotec, 130-097-049) and LS-columns (Miltenyi Biotec, 130-042-401). Separated populations were stained with PI and CD133-APC conjugated antibody. Cells sorting was performed using BD LSRII flow cytometer using Diva software (BD Biosciences). Cells were collected and pelleted for RNAseq and mass spectrometry analysis.

Separations were performed on Control and EIF4G2KD HEC-1A cells using magnetic beads for CD133 (CD133 Microbead Kit, Miltenyi Biotec, 130-097-049), or CD44 (CD44 Microbeads Kit, Miltenyi Biotec, 130-095-194) for RL95-2 cells, on LS-columns (Miltenyi Biotec, 130-042-401) according to the manufacturer’s recommendations.

### MARS-seq library preparation and data analysis

15,000 CD133+ were FACS sorted into 40 μL of lysis binding buffer and mRNA was isolated using Dynabeads mRNA DIRECT Purification Kit (Thermo-Fisher scientific; cat# 61012) according to manufacturer’s protocol. MARSseq Libraries were generated by the MARS-seq protocol (46). The analysis of the MARS-seq data with the UTAP (User-friendly Transcriptome Analysis) Pipeline, alignment to hg38 with STAR v2.4.2a, normalization and differential expression analysis by DESeq2 were all performed as previously described (47). Raw P values were adjusted for multiple testing using the procedure of Benjamini and Hochberg. Genes with baseMean expression > 10, log2FC > 0.58 (fold change > 1.5), and p.adj < 0.05 were considered as genes with differential expression.

### Mass spectrometry

Sorted cell pellets were lysed in 50mM Tris-HCl pH 7.4, 5% SDS, and sonicated (Bioruptor Pico, Diagenode, USA). Protein concentration was measured using the BCA assay (Thermo Scientific, USA). From each sample, 20μg of total protein was subjected to in-solution tryptic digestion using the suspension trapping (S-trap) method as previously described (48). Liquid chromatography and Mass Spectrometry was performed as described previously (BioRxivs 2023.08.22.554280), using split-less nano-Ultra Performance Liquid Chromatography, reversed-phase Symmetry C18 trapping column (Waters) and a T3 HSS nano-column (Waters) for desalting and separation, coupled to a quadrupole orbitrap mass spectrometer (Q Exactive HFX, Thermo Scientific). Data acquisition, processing and analysis was performed as described (21). Differential protein abundance was tested using the glht R function by ANOVA followed by Tukey analysis on the intensity values using a logarithmic scale. Changes in protein abundance of at least 1.5 between conditions, with *p*.value < 0.05 were considered significant.

### Cell viability assay

Cells were plated in sealed white 96-well plates (5000 cells/well). Luminescence was measured 24 h after plating (T0) and after 4 d using the CellTiter-Glo Luminescent Cell Viability Assay protocol (Promega; cat# G7573). When indicated, cells were treated 24 h after plating with 2.5 nM Taxol (Paclitaxel; Sigma-Aldrich; cat# 7402) or subjected to 8 or 16 Gy X-Ray Irradiation using XRAD 320 (Precision X-Ray). Control cells were subjected to incubation with DMSO or mock irradiation, respectively. CellTiter-Glo luminesce values of treated cells were normalized to that of control cells 4 d after treatment to obtain the relative viability.

### Western blot analysis

Cells were lysed in RIPA lysis buffer (20 mM Tris, pH 8.5, 0.1% NP40, 150mM NaCl, 0.5% sodium deoxycholate, 0.1% SDS) supplemented with 10 µl/ml 0.1M PMSF (Sigma-Aldrich, 93482), 1% protease inhibitor (Sigma-Aldrich, P8340) and 1% phosphatase inhibitor (Sigma-Aldrich; P5726). Proteins were separated by SDS-PAGE and transferred to nitrocellulose membrane or PVDF, which were incubated with the following antibodies: mouse anti-EIF4G2 (BD Biosciences Cat# 610742, RRID:AB_398065), rabbit anti-KLC1 (Abcam Cat# ab174273, RRID:AB_2783556), rabbit anti-KIF5B (Abcam Cat# ab167429, RRID:AB_2715530), rabbit anti-KLC2 (Abcam, ab254848), mouse anti-ALDH1A (Thermo Fisher Scientific Cat# MA5-29023, RRID:AB_2784960) and rabbit anti-γH2AX (Cell Signaling Technology Cat# 2577, RRID:AB_2118010), all diluted 1:1000; mouse anti-GAPDH (Millipore Cat# MAB374, RRID:AB_2107445), diluted 1:3000, and mouse anti-Tubulin (Sigma-Aldrich Cat# T9026, RRID:AB_477593), diluted 1:70000. Detection was done with either horseradish peroxidase (HRP)-conjugated goat anti-mouse (Jackson ImmunoResearch Labs Cat# 115-035-003, RRID: AB_10015289) or anti-rabbit (Jackson ImmunoResearch, 111-165-144) secondary antibodies, followed by enhanced chemiluminescence using EZ-ECL (Biological Industries, 20-500). Analysis of western blot results was done using ImageJ software. Quantification of protein steady levels was performed and normalized to the appropriate loading control and log transformed.

### Polysomal profiling

Control and EIF4G2KD HEC-1A CD133+ and DC133-fractionated populations were subjected to polysomal fractionation as described (15).

### Immunofluorescent staining of human tumor microarray

Human tumor microarrays (TMA) containing samples from patients were deparaffinized and fixed with 10% neutral buffered formalin. Antigen retrieval was performed using citrate buffer (pH 6.0). Slides were then blocked with 10% BSA + 0.05% Tween20 and the antibodies were diluted in 2% BSA in 0.05% PBST and used in a multiplexed manner with OPAL reagents (Akoya Biosciences). All primary antibodies (described above for western blotting) were incubated overnight at 4°C (1:400) according to the following staining sequences: Set 1: KLC1 → KIF5B → CK → DAPI; Set 2: EIF4G2 → CK → DAPI. Each antibody was validated and optimized separately and then the multiplex staining was optimized. Briefly, following primary antibody incubation, slides were washed with 0.05% PBST, incubated with secondary antibodies conjugated to HRP (1:400), washed again, and incubated with OPAL reagents. Slides were then washed, and antigen retrieval was performed. Then, slides were washed with PBS and stained with the next primary antibody or with DAPI at the end of the cycle. Finally, slides were mounted using Immu-mount (#9990402, Thermo Scientific). Images were taken with a Phenocycler scanner (Akoya®) and analyzed using QuPath software. Cell segmentation was done using Cellpose. For each patient, average cell intensities for the indicated markers was calculated, and correlation between the different markers was analyzed.

### Survival analysis

Patients were stratified based on different parameters (staining intensities, tumor grade, stage and type). Kaplan Meier (KM) analysis of overall survival with log rank P value was performed on patients stratified by median expression of each of these signatures. Paired comparisons were calculated using the Survdiff function in R with FDR correction.

### Statistical analysis

All additional statistical analyses not described above were performed using Prism 9.3 software (GraphPad Software), as specified in figure legends. Non-significant comparisons were mostly not marked in the graphs.

### Data availability

The mass spectrometry proteomics data have been deposited in the MassIve repository of the ProteomeXchange consortium (https://massive.ucsd.edu), with the dataset identifier MSV000092778. The RNA-seq data generated in this study is publicly available in Gene Expression Omnibus (GEO) at GSE242717.

## Acknowledgements

We would like to thank Dr. Anastasia Lev and Prof. Rivka Dikstein from the Weizmann Institute for the polysomal profile analysis. This research was supported by grants from the Women’s Health Research Center at the Weizmann and the Pasteur-Weizmann Council for A.K, and by a scholarship from Emek Medical Center for M.M.A.

## References

1. de la Parra C, Walters BA, Geter P, Schneider RJ. Translation initiation factors and their relevance in cancer. Curr Opin Genet Dev 2018;48:82–8

2. Imataka H, Olsen HS, Sonenberg N. A new translational regulator with homology to eukaryotic translation initiation factor 4G. EMBO J 1997;16:817–25

3. Levy-Strumpf N, Deiss LP, Berissi H, Kimchi A. DAP-5, a novel homolog of eukaryotic translation initiation factor 4G isolated as a putative modulator of gamma interferon-induced programmed cell death. Mol Cell Biol 1997;17:1615–25

4. Shaughnessy JD, Jenkins NA, Copeland NG. cDNA cloning, expression analysis, and chromosomal localization of a gene with high homology to wheat eif-(iso)4F and mammalian eIF-4G. Genomics 1997;39:192–7

5. Yamanaka S, Poksay KS, Arnold KS, Innerarity TL. A novel translational repressor mRNA is edited extensively in livers containing tumors caused by the transgene expression of the apoB mRNA-editing enzyme. Genes Dev 1997;11:321–33

6. Liberman N, Gandin V, Svitkin YV, David M, Virgili G, Jaramillo M, et al. DAP5 associates with eIF2beta and eIF4AI to promote Internal Ribosome Entry Site driven translation. Nucleic Acids Res 2015;43:3764–75

7. de la Parra C, Ernlund A, Alard A, Ruggles K, Ueberheide B, Schneider RJ. A widespread alternate form of cap-dependent mRNA translation initiation. Nat Commun 2018;9:3068

8. Smirnova VV, Shestakova ED, Nogina DS, Mishchenko PA, Prikazchikova TA, Zatsepin TS, et al. Ribosomal leaky scanning through a translated uORF requires eIF4G2. Nucleic Acids Res 2022

9. Weber R, Kleemann L, Hirschberg I, Chung MY, Valkov E, Igreja C. DAP5 enables main ORF translation on mRNAs with structured and uORF-containing 5’ leaders. Nat Commun 2022;13:7510

10. Yang Y, Fan X, Mao M, Song X, Wu P, Zhang Y, et al. Extensive translation of circular RNAs driven by N(6)-methyladenosine. Cell Res 2017;27:626–41

11. Yamanaka S, Zhang XY, Maeda M, Miura K, Wang S, Farese RV, Jr., et al. Essential role of NAT1/p97/DAP5 in embryonic differentiation and the retinoic acid pathway. EMBO J 2000;19:5533–41

12. Yoshikane N, Nakamura N, Ueda R, Ueno N, Yamanaka S, Nakamura M. Drosophila NAT1, a homolog of the vertebrate translational regulator NAT1/DAP5/p97, is required for embryonic germband extension and metamorphosis. Dev Growth Differ 2007;49:623–34

13. Nousch M, Reed V, Bryson-Richardson RJ, Currie PD, Preiss T. The eIF4G-homolog p97 can activate translation independent of caspase cleavage. RNA 2007;13:374–84

14. Sugiyama H, Takahashi K, Yamamoto T, Iwasaki M, Narita M, Nakamura M, et al. Nat1 promotes translation of specific proteins that induce differentiation of mouse embryonic stem cells. Proc Natl Acad Sci U S A 2017;114:340–5

15. Yoffe Y, David M, Kalaora R, Povodovski L, Friedlander G, Feldmesser E, et al. Cap-independent translation by DAP5 controls cell fate decisions in human embryonic stem cells. Genes Dev 2016;30:1991–2004

16. Warnakulasuriyarachchi D, Cerquozzi S, Cheung HH, Holcik M. Translational induction of the inhibitor of apoptosis protein HIAP2 during endoplasmic reticulum stress attenuates cell death and is mediated via an inducible internal ribosome entry site element. J Biol Chem 2004;279:17148–57

17. Henis-Korenblit S, Shani G, Sines T, Marash L, Shohat G, Kimchi A. The caspase-cleaved DAP5 protein supports internal ribosome entry site-mediated translation of death proteins. Proceedings of the National Academy of Sciences of the United States of America 2002;99:5400–5

18. Hundsdoerfer P, Thoma C, Hentze MW. Eukaryotic translation initiation factor 4GI and p97 promote cellular internal ribosome entry sequence-driven translation. Proc Natl Acad Sci U S A 2005;102:13421–6

19. Marash L, Liberman N, Henis-Korenblit S, Sivan G, Reem E, Elroy-Stein O, et al. DAP5 promotes cap-independent translation of Bcl-2 and CDK1 to facilitate cell survival during mitosis. Mol Cell 2008;30:447–59

20. Weingarten-Gabbay S, Khan D, Liberman N, Yoffe Y, Bialik S, Das S, et al.The translation initiation factor DAP5 promotes IRES-driven translation of p53 mRNA. Oncogene 2014;33:611–8

21. David M, Olender T, Mizrahi O, Weingarten-Gabbay S, Friedlander G, Meril S, et al. DAP5 drives translation of specific mRNA targets with upstream ORFs in human embryonic stem cells. RNA 2022;28:1325–36

22. Alard A, Katsara O, Rios-Fuller T, Parra C, Ozerdem U, Ernlund A, et al. Breast cancer cell mesenchymal transition and metastasis directed by DAP5/eIF3d-mediated selective mRNA translation. Cell Rep 2023;42:112646

23. Fu L, Wang Z, Jiang F, Wei G, Sun L, Guo C, et al. High Expression of EIF4G2 Mediated by the TUG1/Hsa-miR-26a-5p Axis Is Associated with Poor Prognosis and Immune Infiltration of Gastric Cancer. J Oncol 2022;2022:9342283

24. Buim ME, Soares FA, Sarkis AS, Nagai MA. The transcripts of SFRP1,CEP63 and EIF4G2 genes are frequently downregulated in transitional cell carcinomas of the bladder. Oncology 2005;69:445–54

25. Siegel RL, Miller KD, Fuchs HE, Jemal A. Cancer statistics, 2022. CA: A Cancer Journal for Clinicians 2022;72:7–33

26. Makker V, MacKay H, Ray-Coquard I, Levine DA, Westin SN, Aoki D, et al. Endometrial cancer. Nat Rev Dis Primers 2021;7:88

27. Giannone G, Attademo L, Scotto G, Genta S, Ghisoni E, Tuninetti V, et al. Endometrial Cancer Stem Cells: Role, Characterization and Therapeutic Implications. Cancers (Basel) 2019;11

28. Banz-Jansen C, Helweg LP, Kaltschmidt B. Endometrial Cancer Stem Cells: Where Do We Stand and Where Should We Go? Int J Mol Sci 2022;23

29. Kuramoto H. Studies of the growth and cytogenetic properties of human endometrial adenocarcinoma in culture and its development into an established line. Acta Obstet Gynaecol Jpn 1972;19:47–58

30. Lelle RJ, Talavera F, Gretz H, Roberts JA, Menon KM. Epidermal growth factor receptor expression in three different human endometrial cancer cell lines. Cancer 1993;72:519–25

31. Way DL, Grosso DS, Davis JR, Surwit EA, Christian CD. Characterization of a new human endometrial carcinoma (RL95-2) established in tissue culture. In Vitro 1983;19:147–58

32. Lu KH, Broaddus RR. Endometrial cancer. N Engl J Med 2021:181–93

33. Ben-Ari Fuchs S, Lieder I, Stelzer G, Mazor Y, Buzhor E, Kaplan S, et al. GeneAnalytics: An Integrative Gene Set Analysis Tool for Next Generation Sequencing, RNAseq and Microarray Data. OMICS 2016;20:139–51

34. Yue H, Hu Z, Hu R, Guo Z, Zheng Y, Wang Y, et al. ALDH1A1 in Cancers: Bidirectional Function, Drug Resistance, and Regulatory Mechanism. Front Oncol 2022;12:918778

35. Mori Y, Yamawaki K, Ishiguro T, Yoshihara K, Ueda H, Sato A, et al. ALDH-Dependent Glycolytic Activation Mediates Stemness and Paclitaxel Resistance in Patient-Derived Spheroid Models of Uterine Endometrial Cancer. Stem Cell Reports 2019;13:730–46

36. Chen G, Liu B, Yin S, Li S, Guo Ye, Wang M, et al. Hypoxia induces an endometrial cancer stem-like cell phenotype via HIF-dependent demethylation of SOX2 mRNA. Oncogenesis 2020;9

37. Kiyohara MH, Dillard C, Tsui J, Kim SR, Lu J, Sachdev D, et al. EMP2 is a novel therapeutic target for endometrial cancer stem cells. Oncogene 2017;36:5793–807

38. Silva IA, Bai S, McLean K, Yang K, Griffith K, Thomas D, et al. Aldehyde dehydrogenase in combination with CD133 defines angiogenic ovarian cancer stem cells that portend poor patient survival. Cancer Research 2011;71:3991–4001

39. Nishimura K, Fukuda A, Hisatake K. Mechanisms of the Metabolic Shift during Somatic Cell Reprogramming. Int J Mol Sci 2019;20

40. Zhu X, Chen HH, Gao CY, Zhang XX, Jiang JX, Zhang Y, et al. Energy metabolism in cancer stem cells. World J Stem Cells 2020;12:448–61

41. Rath O, Kozielski F. Kinesins and cancer. Nat Rev Cancer 2012;12:527–39

42. Lottersberger F, Karssemeijer RA, Dimitrova N, de Lange T. 53BP1 and the LINC Complex Promote Microtubule-Dependent DSB Mobility and DNA Repair. Cell 2015;163:880–93

43. Moamer A, Hachim IY, Binothman N, Wang N, Lebrun JJ, Ali S. A role for kinesin-1 subunits KIF5B/KLC1 in regulating epithelial mesenchymal plasticity in breast tumorigenesis. EBioMedicine 2019;45:92–107

44. Marchesin V, Castro-Castro A, Lodillinsky C, Castagnino A, Cyrta J, Bonsang-Kitzis H, et al. ARF6-JIP3/4 regulate endosomal tubules for MT1-MMP exocytosis in cancer invasion. J Cell Biol 2015;211:339–58

45. Cui J, Jin G, Yu B, Wang Z, Lin R, Huang J-D. Stable knockdown of Kif5b in MDCK cells leads to epithelial–mesenchymal transition. Biochemical and Biophysical Research Communications 2015;463:123–9

46. Keren-Shaul H, Kenigsberg E, Jaitin DA, David E, Paul F, Tanay A, et al. MARS-seq2.0: an experimental and analytical pipeline for indexed sorting combined with single-cell RNA sequencing. Nat Protoc 2019;14:1841–62

47. Parichha A, Suresh V, Chatterjee M, Kshirsagar A, Ben-Reuven L, Olender T, et al. Constitutive activation of canonical Wnt signaling disrupts choroid plexus epithelial fate. Nat Commun 2022;13:633

48. Elinger D, Gabashvili A, Levin Y. Suspension Trapping (S-Trap) Is Compatible with Typical Protein Extraction Buffers and Detergents for Bottom-Up Proteomics. J Proteome Res 2019;18:1441–5

