## Supplementary Figures for "Loss of EIF4G2 Mediates Aggressiveness in Distinct Human Endometrial Cancer Subpopulations with Poorer Survival Outcome in Patients"

**Meril, et al. Supplementary Information- Figures and Tables**

### Supplementary Table S1- Patient information (excel table)

Details on patient information, diagnosis, tumor histology, therapies received, recurrence and survival data. NA, data not available. 0 and 1 are indication for "No" and "Yes" respectively. Also indicated are the staining intensities for KLC1, KIF5B and EIF4G2 proteins in each corresponding patient's tumor core. High and low staining intensity is determined by values above and below the median staining intensity for each protein: KLC1, 55.4522; KIF5B, 15.846; EIF4G2, 3.7002.

### Supplementary Table S2- RNA-seq data (excel table)

Normalized data from RNA-seq analysis is shown for all genes detected by MARS-Seq for comparisons among sorted CD133- and CD133+ control and EIF4G2 KD HEC-1A cells (first tab). Remaining tabs show the significant DEGs in the individual comparisons, sorted from lowest to highest log2 fold-change value. Negative fold-changes represent genes with decreased expression in the EIF4G2 KD vs control comparisons (second and third tabs) or the CD133- vs CD133+ populations (third and fourth tabs).

### Supplementary Table S3- Mass Spectrometry data (excel table)

Normalized data from MS analysis is shown for all detected proteins for comparisons among sorted CD133- and CD133+ control and EIF4G2 KD HEC-1A cells (first tab). Remaining tabs show the proteins with significant changes in abundance in the individual comparisons, sorted from lowest to highest log2 fold-change value. Proteins with lower abundance in the EIF4G2 KD (CD133- or CD133+) cells compared to control KD cells are highlighted in orange, and greater abundance in blue (second and third tabs). Proteins with lower abundance in the CD133- cells (control or EIF4G2 KD) compared to CD133+ cells are highlighted in orange, and greater abundance in blue (third and fourth tabs).

### Meril et al. Supplementary Fig. S1

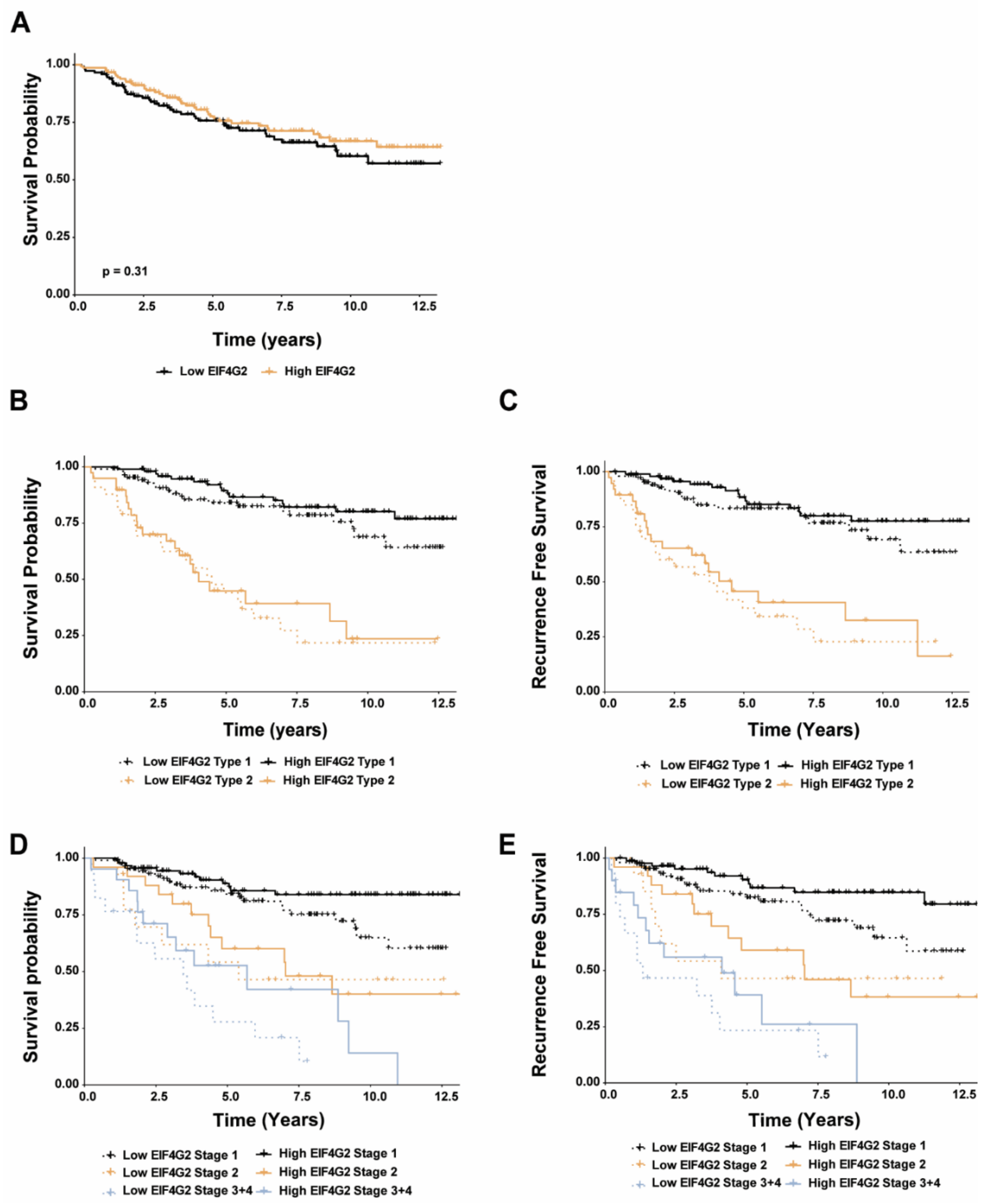


**Supplementary Fig. S1- Additional survival analysis of EC patients according to EIF4G2 expression levels.**

FFPE TMA sections from the 280 patients were immunostained for EIF4G2, CK and DAPI. **A-D** Patients were stratified according to high and low intensity levels of EIF4G2 staining in CK positive cells, and overall survival (**A, B, D)** or recurrence free survival (**C, E**) was assessed by Kaplan-Meier analysis in all patients **(A)** or in patients stratified according to tumor type (**B, C)** and stage **(D, E)**. Log rank *P*-val was determined on patients stratified by median expression of EIF4G2 intensity. Paired comparison was calculated with FDR correction.

Meril et al. Supplementary Fig. S2

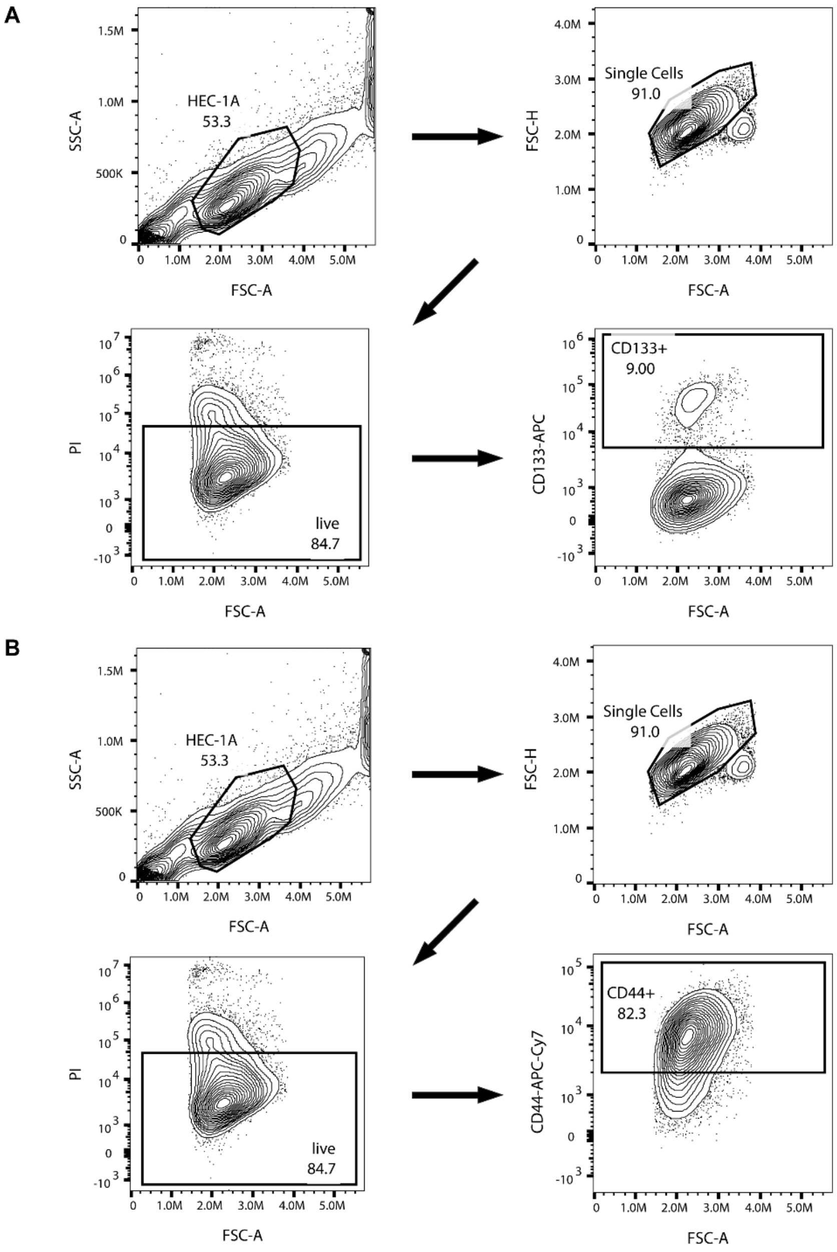


**Supplementary Fig. S2- Strategies for FACS sorting by CD133 and CD44 expression. A, B** Gating strategies for FACS analysis and sorting for CD133 **(A)** and CD44 **(B**) in HEC-1A cells. In each step, the boxed cells were taken for further analysis or sorting in the next step. Numbers represent total percentages of cells within the boxed population.

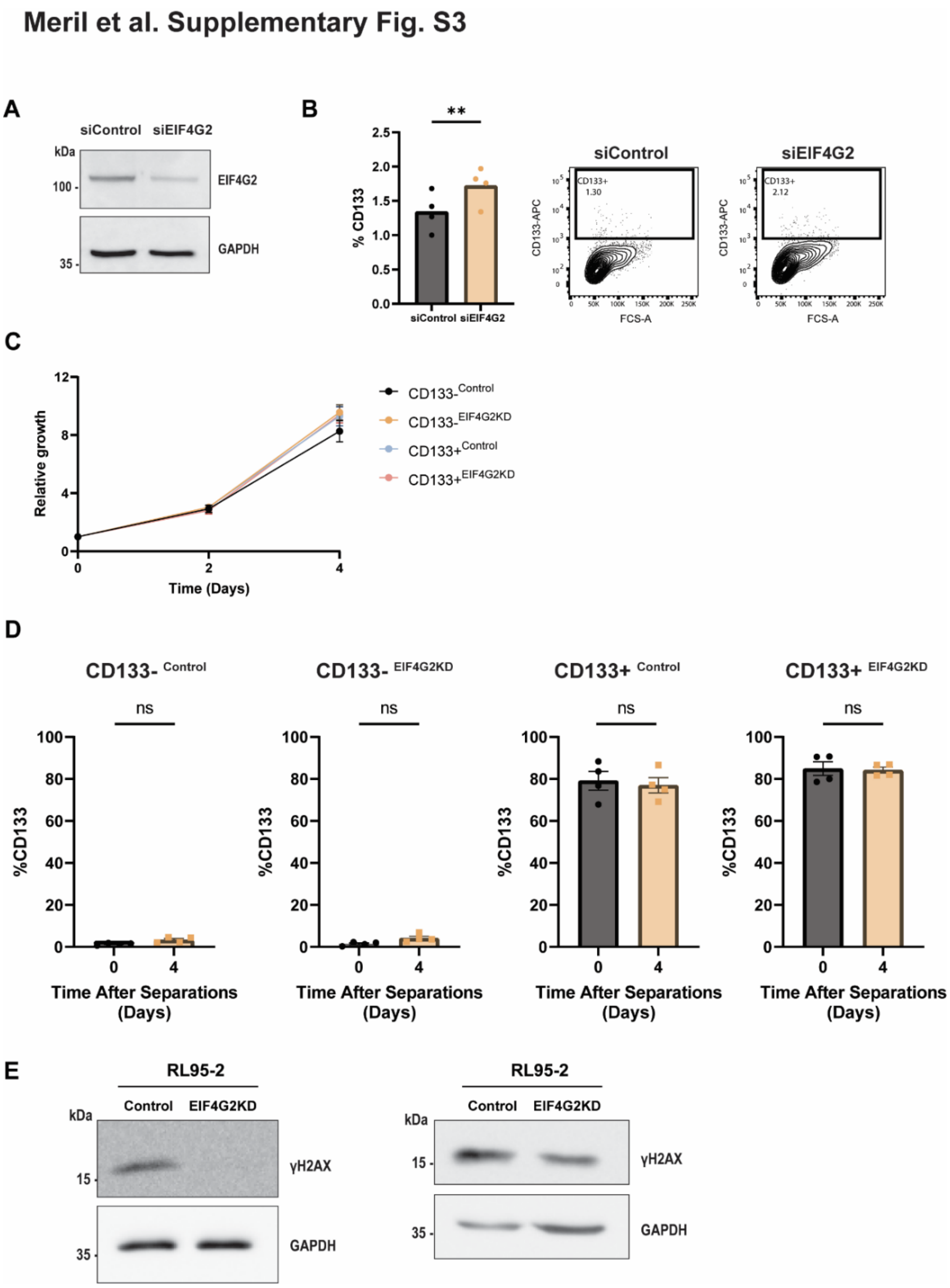


**Supplementary Fig. S3- Additional analysis of EIF4G2 KD EC cells.**

**A, B** HEC-1A cells were transfected with control siRNA or siRNA against EIF4G2 to generate transient KD cells. **(A)** Total cell extract was subjected to western bot analysis for EIF4G2 or GAPDH as loading control. Representative blot of n=5 independent experiments is shown. **(B)** Flow cytometry analysis for CD133 was performed on siRNA transfected cells. Quantification of percent positive cells is presented as mean with individual points plotted, n=4 independent experiments. *P*-val was calculated using two-tailed t-test (**: $p$<0.01). A representative flow cytometry plot is shown. **C** Relative growth of CD133- and CD133+ separated control and EIF4G2

KD HEC-1A cells on days 0, 2 and 4 as determined by CellTiter-Glo assay. Luminescence values were normalized to Day 0 of the same cell type. Shown are mean values±SEM of n=6 independent experiments. **D** Flow cytometry analysis for CD133 expression over 4 d in CD133- and CD133+ separated control and EIF4G2 KD HEC-1A cells. Statistical analysis was determined using one-way ANOVA relative to day 0. Shown are mean values±SEM of n=3 independent experiments. **E** Additional representative blots of cell lysates form RL95-2 control and EIF4G2 KD cells that were subjected to western blot analysis for γH2AX and GAPDH antibodies as loading control 4 d following irradiation.

Meril et al. Supplementary Fig. S4

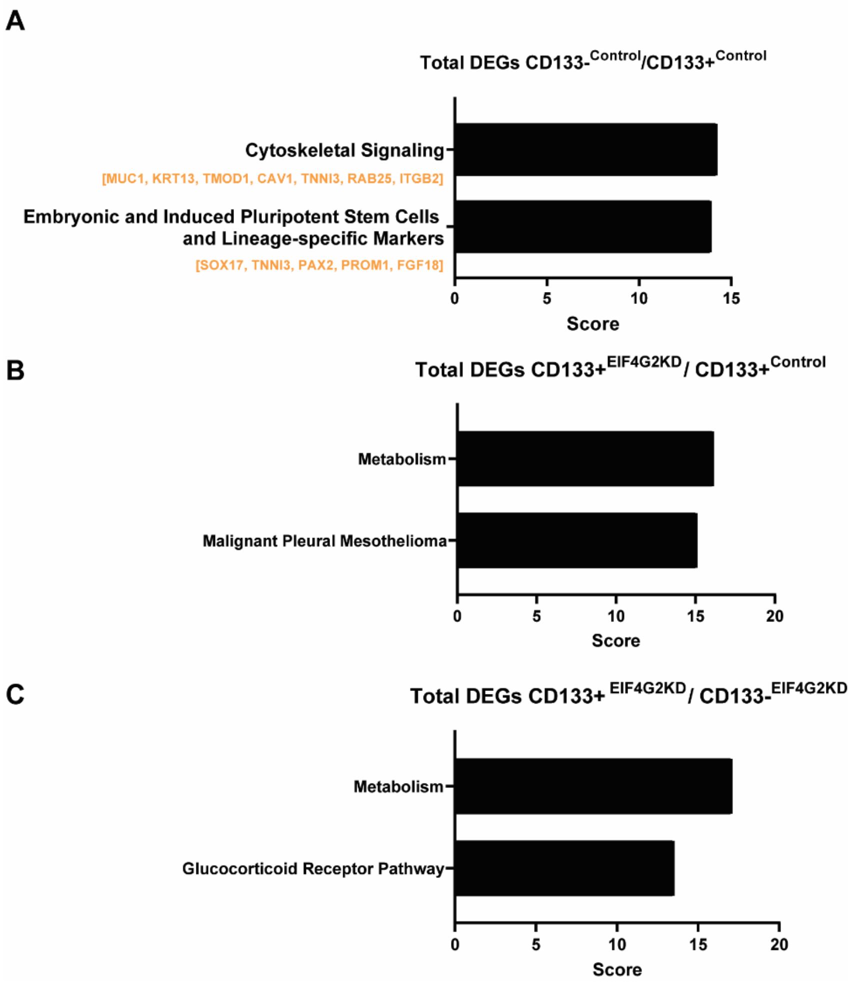


**Supplementary Fig. S4- Pathway analysis of RNA-seq results in sorted EIF4G2 KD cells**

**A-C** RNA-seq analysis was performed on control and EIF4G2 HEC-1A cells that were FACS sorted for CD133 surface expression. Pathway analysis of paired comparisons was conducted by GeneAnalytics. Only high score pathways are shown for **(A)** Total DEGs of the $CD133^{-Control}/CD133^{+Control}$ comparison, **(B)** total DEGs of the $CD133^{+EIF4G2KD}/CD133^{+Control}$ comparison or **(C)** total DEGs of the $CD133^{+EIF4G2KD}/CD133^{-EIF4G2}$ comparison.

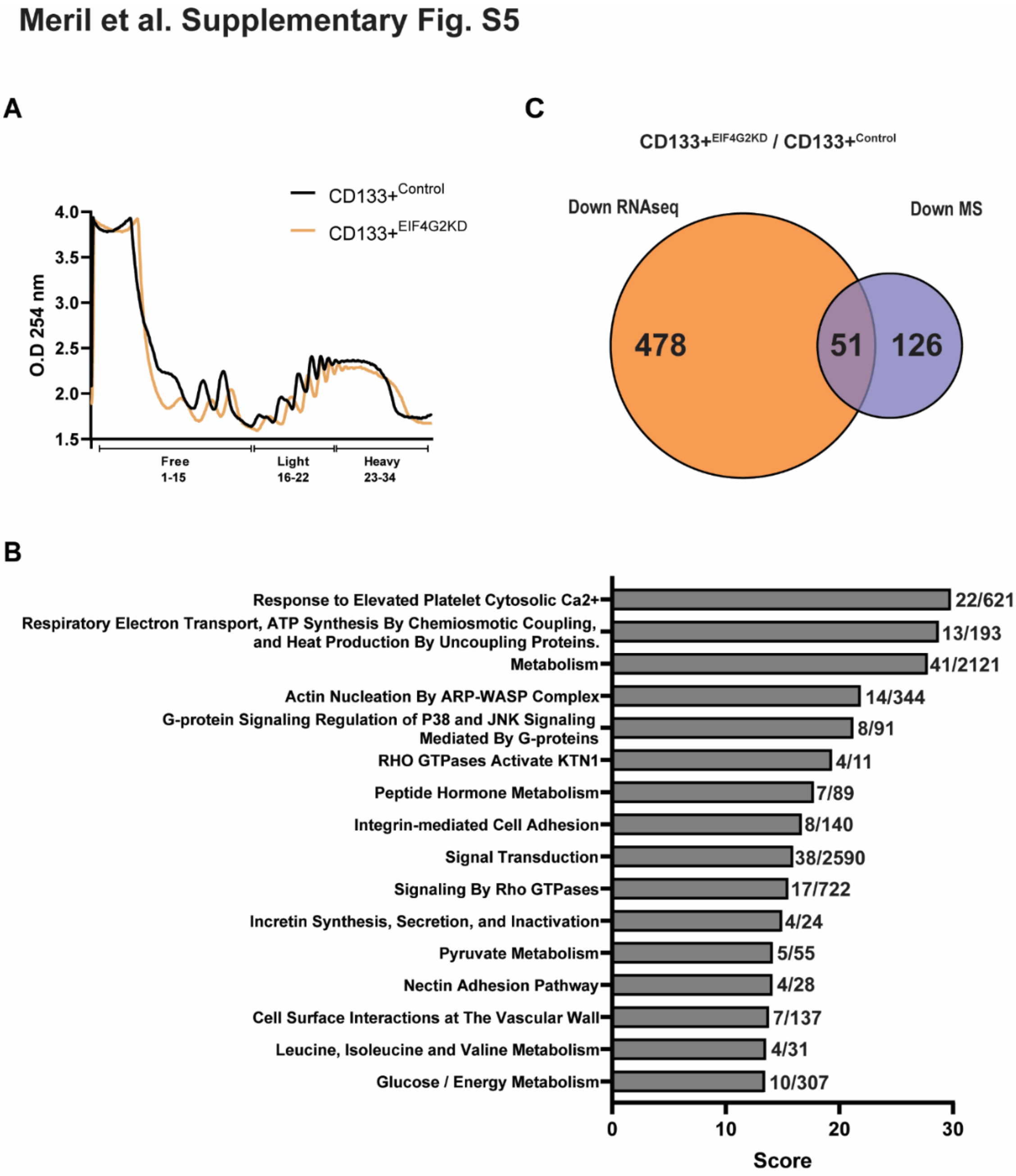


**Supplementary Fig. S5- Additional proteomic analysis of sorted EIF4G2 KD cells.**

**A** Polysomal profile of CD133+ control and EIF4G2 KD HEC-1A cells. Values are presented as O.D. at 250nm. Fractions are listed on X-axis, representing free ribosomes, light and heavy polysomes as indicated. A representative plot of n=3 independent experiments is shown. **B** High scoring significant pathways identified by GeneAnalytics pathway analysis of the set of proteins with decreased abundance in the $CD133+^{EIF4G2KD}$ / $CD133+^{Control}$ comparison. Score numbers indicating significance are indicated. Only high score pathways are presented. Numbers at right represent the number of proteins identified in the dataset out of the total number of proteins within the given pathway. **C** Venn diagram showing overlap between downregulated DEGs as identified in the RNA-seq analysis and proteins with decreased abundance in MS in the comparison of $CD133+^{EIF4G2KD}$/$CD133+^{Control}$ populations.

### Meril et al. Supplementary Fig. S6

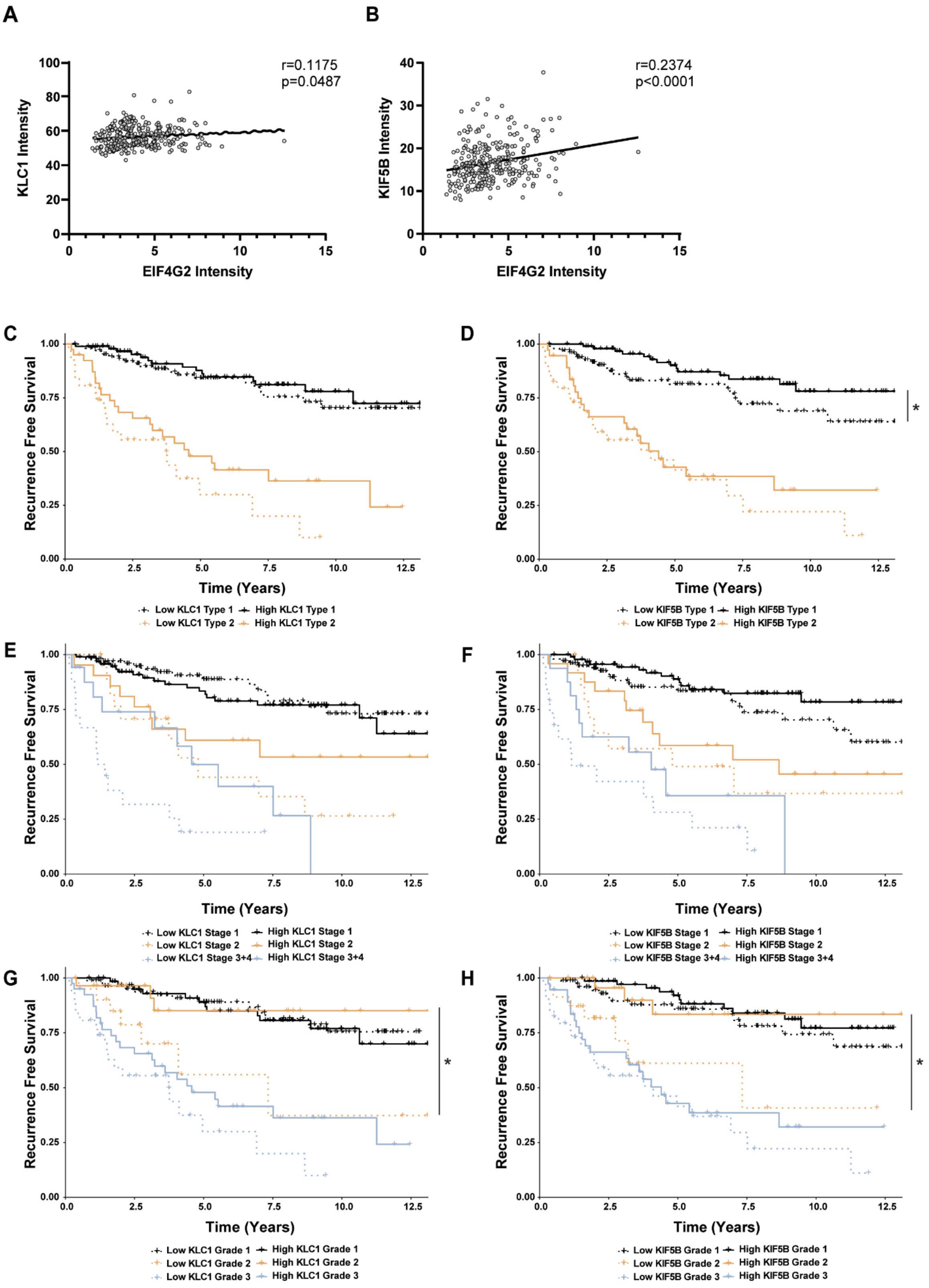


**Supplementary Fig. S6- Additional analysis of KLC1 and KIF5B expression in EC patients and association with recurrence-free survival.**

FFPE TMA sections from the cohort of 280 patients were immunostained for EIF4G2 and CK in the first panel or KLC1, KIF5B and CK in the second panel. **A, B** correlation analysis between KLC1 and EIF4G2 **(A)** or KIF5B and EIF4G2 **(B)** staining intensities was performed. Correlation coefficient r and *P*-val are noted on the graph. **C-H** Recurrent free survival of 280 endometrial patients according to tumor **(C, D)** type, **(E, F)** stage and **(G, H)** grade. KLC1 **(C, E, G)** and KIF5B **(D, F, H)** staining in CK positive cells was stratified according to high and low intensity levels compared to the calculated median and recurrent free survival was assessed by Kaplan-Meier statistics. Log rank *P*-value was determined for protein expression of each of these signatures. Paired comparison was calculated with FDR correction. *: $p<0.05$.
